# Estimating the effect size of a hidden causal factor between SNPs and a continuous trait: a mediation model approach

**DOI:** 10.1101/2022.06.28.497617

**Authors:** Zhuoran Ding, Marylyn D. Ritchie, Benjamin F. Voight, Wei-Ting Hwang

## Abstract

**Background:** Observational studies and Mendelian randomization experiments have been used to identify many causal factors for complex traits in humans. Given a set of causal factors, it is important to understand the extent to which these causal factors explain some, all, or none of the genetic heritability, as measured by single-nucleotide polymorphisms (SNPs) that are associated with the trait. Using the mediation model framework with SNPs as the exposure, a trait of interest as the outcome, and the known causal factors as the mediators, we hypothesize that any unexplained association between the SNPs and the outcome trait is mediated by an additional unobserved, hidden causal factor.

**Results:** We propose a method to infer the effect size of this hidden mediating causal factor on the outcome trait by utilizing the estimated associations between a continuous outcome trait, the known causal factors, and the SNPs. The proposed method consists of three steps and, in the end, implements Markov chain Monte Carlo to obtain a posterior distribution for the hidden mediator’s effect size. We evaluate our proposed method via extensive simulations and show that when the model assumptions hold, our method estimates the effect size of the hidden mediator well and controls type I error rate if the hidden mediator does not exist.. In addition, we apply the method to the UK Biobank data and find that a potential hidden mediator for waist-hip ratio exists in the European population, and the hidden mediator has a large effect size relatively to the effect size of the known mediator BMI.

**Conclusions:** We develop a framework to infer the effect of potential, hidden mediators influencing complex traits. This framework can begin to place boundaries on unexplained risk factors contributing to complex traits.

## 1 Introduction

One of the goals in studying the associations between heritable traits and disease outcomes is to identify which of these factors are truly causal for the outcome. Expansion of causal inference studies like Mendelian randomization studies [1] in recent years have provided one piece of causal evidence to risk factors identified from observational epidemiological studies [2]. However, generating causal evidence between exposures and disease outcome leaves unaddressed the question of whether the genetic heritability of the trait can be fully explained by the set of known causal factors or if there exist additional ones that are unidentified but further explain disease risk or trait heritability. For example, it is known that high body mass index (BMI) and potentially low-density lipoprotein cholesterol (LDLC) increase the risk of developing type 2 diabetes (T2D) [3,4], but can the genetic heritability of T2D be fully explained by BMI and LDL-C? If the heritability of a disease can be fully explained by the set of existing known causal factors, then research on the disease can focus on studying the biological mechanisms of those causal factors. On the other hand, if there is genetic heritability that remains unexplained, other causal factors for the disease may exist and are currently hidden from observation. Studying the characteristics of the hidden causal factors may provide insights into the novel biological associations and mechanisms that remain undiscovered.

The classic mediation framework decomposes the associations between the exposures and the outcome into direct effect and indirect effect through a mediator (i.e., mediated effects) [5, 6]. The question described earlier can be considered within a mediation framework with the trait of interest as the outcome, identified single nucleotide polymorphisms (SNPs) associated with the trait as the exposure, and the known causal factors of the trait as the mediators. Under this framework, any remaining direct effects between the identified SNPs and the outcome trait are viewed as the residual associations that are not explained by causal factors or mediators included in the model. Thus, the residual associations could be due to one or more hidden mediators. As the first step towards learning about the residual associations, in what follows we consider the simplest case that the remaining direct effects between the identified SNPs and the outcome trait are due to a single hidden mediator. Under this case, we can further decompose the unexplained genetic heritability into two parts: (i) the SNP effects on the hidden mediator, and (ii) the effect size of the hidden mediator on the outcome trait. The goal of this study is to estimate the latter: the effect size of the hidden mediator on the outcome trait.

In this work, we assume that the values of the SNPs, the known mediators, and the outcome trait are accurately measured and there are no unmeasured confounding variables in the model. Furthermore, we assume that the effect sizes of the SNPs on the standardized known and hidden mediators have the same distribution. This is reasonable due to the following: because there is little or no prior information about the hidden mediator, the unexplained genetic heritability can theoretically be decomposed by infinitely many combinations of the two parts. Thus, to limit the number of possible combinations of the two parts and infer a reasonable range of the hidden mediator’s effect size on the outcome, we propose to restrict the effect sizes of the SNPs on the hidden mediator to be similar to the effect sizes of the SNPs on the known mediators. Under this assumption, one part of the unexplained genetic heritability, that is, the SNP effects on the hidden mediator, can be learned from the SNP effects on the known mediators. Furthermore, the effect size of the hidden mediator on the outcome trait can be inferred by dividing the direct effects between the SNPs and the outcome trait by the inferred SNP effects on the hidden mediator.

The rest of the report is organized as follows: In Section 2, we provide a broad overview of our approach and describe each step of our method in detail. We also describe the settings for the simulation study. In Section 3, we present the simulation results and the application on investigating the trait of waist-hip ratio. We conclude in Section 4 with a discussion.

## 2 Methods

### 2.1 The mediation model and notations

We illustrate our method using a model with two known mediators, although our method can extend to the case with more than two mediators or only one known mediator. We denote a continuous trait of interest as *Y*, the vector of SNPs associated with *Y* as ***G***, and the two known mediating causal factors of *Y* as *M*_1_ and *M*_2_ (Figure 1A). *M*_1_ and *M*_2_ are both standardized to have unit variance. Furthermore, the SNP effects from *G* to *M*_1_ are represented by a vector ***a***_1_ and the SNP effects from ***G*** to *M*_2_ are represented by a vector ***a***_2_, the direct effects between ***G*** and *Y* are represented by a vector ***c***, the effect size of *M*_1_ on *Y* is denoted by a scalar *b*_1_, and the effect size of *M*_2_ on *Y* is denoted by a scalar *b*_2_.

**Figure 1:**
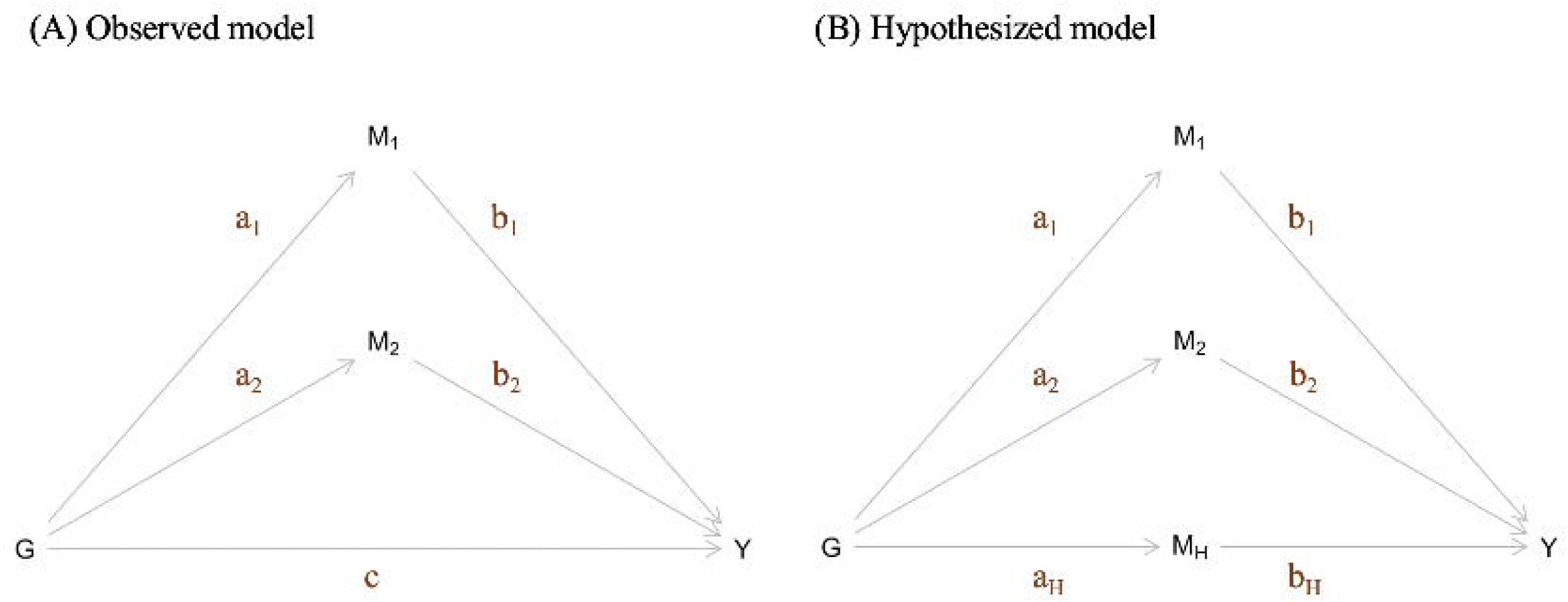
The mediation model framework. (A) the observed model. (B) the hypothesized model.

If there is an unexplained genetic heritability between ***G*** and *Y* (i.e., *c* ≠ 0), then we assume a hidden mediator exists and is denoted by *M*_*H*_ (Figure 1B). The SNP effects from ***G*** to *M*_*H*_ are represented by a vector ***a***_***H***_. The goal of the proposed work is to infer the effect size of the hidden mediator *M*_*H*_ on *Y*, denoted by *b*_*H*_. To account for the scenario that some of the SNPs in ***G*** are not associated with *M*_*H*_, we use *π*_*H*_ to denote the proportion of the SNPs in ***G*** that are associated with *M*_*H*_.

If we were to observe *M*_*H*_, the direct effect vector *c* can be decomposed as ***c* = *a***_***H***_*b*_*H*_. However, since *M*_*H*_ is not observed, we can only estimate ***a***_**1**_, ***a***_**2**_, *b*_1_, *b*_2_, and ***c*** as shown in Figure 1A but not ***a***_***H***_ and *b*_*H*_ as shown in Figure 1B. We propose to use the estimates of ***a***_**1**_, ***a***_**2**_, and ***c***, denoted as ***a***^⍰^_**1**_, ***a***^⍰^_**2**_, and ***c***^⍰^ to infer ***a***_***H***_ and, subsequently, infer *b*_*H*_. For simplicity, we denote the joint vector of ***a***_**1**_ and ***a***_**2**_ and the joint vector of ***a***^⍰^_**1**_ and ***a***^⍰^_**2**_ as ***a*** and ***a***^⍰^, respectively.

### 2.2 Overview of the proposed method and the rationales

To infer the effect size of the hidden mediator, *b*_*H*_, we utilize the fact that we can decompose the direct effect ***c*** as ***a***_***H***_*b*_*H*_. To do so, we assume that the SNP effects on the hidden mediator, ***a***_***H***_, share some similarities with the SNP effects on the known mediators, ***a***_**1**_ and ***a***_**2**_. We first consider the simplest case, in which we assume that all the SNP effects (on both the known and unknown mediators) come from the same distribution. Under this assumption, the true mean and standard deviation of ***a***^⍰^_***H***_ can be consistently estimated by the SNP effects ***a***^⍰^_**1**_ and ***a***^⍰^_**2**_ if the sample size is large and a large number of SNPs are included in the model. We estimate the SNP effects ***a***^⍰^_**1**_ and ***a***^⍰^_**2**_ by fitting two linear regression models with the known mediators *M*_1_ and *M*_2_ as the dependent variables and *G* as the independent variables. Recognizing that assuming similar SNP effect sizes and variances on different mediators may be a strong assumption, we present a more general setting in which the SNP effects could vary according to a three-level structure (Supplementary Materials Section 1) [see Additional file 1].

One challenge in the decomposition of the direct effects of SNPs on Y is that we might not expect every SNP associated with the outcome trait will be associated with the known and hidden mediators. Therefore, we model the SNP effects on the known mediators (***a***), using a mixture model with a point mass at zero and a true effect size distribution that centers at a non-zero value. Because we estimate the SNPs effects on the known mediators (***a***^⍰^) using linear regression models that come with estimation uncertainty, the distribution of ***a***^⍰^ will be a mixture of a distribution centered at zero and the true effect distribution with a non-zero mean and a variance that is larger than the true dispersion. Similarly, the SNP effects on the hidden mediator, ***a***_***H***_, can also be modeled using a mixture model with a point mass at zero and a true effect distribution that centers at a non-zero value. For the same reason as in the case of ***a***^⍰^, the estimated ***c***^⍰^ will be the mixture of a point mass at zero and a true effect distribution not centered at zero and with a larger dispersion. Therefore, we utilize Gaussian mixture models (GMMs) to model the distributions of ***a***^⍰^ and ***c***^⍰^.

### 2.3 The proposed multi-step method

Our method consists of three major steps as shown in Figure 2. In Step 1, we estimate the individual SNP effects on the known mediators to obtain ***a***^⍰^ and estimate the direct effects between the SNPs and the outcome trait to obtain ***c***^⍰^ by fitting a series of linear regression models. In Step 2, we fit GMMs on ***a***^⍰^ and ***c***^⍰^ using an EM algorithm to separate the SNP effects on the known and hidden mediators from the zero-mean noises and estimate the GMM parameters from both distributions. In Step 3, we incorporate the estimated GMM parameters from Step 2 to a GMM Markov chain Monte Carlo (MCMC) procedure to generate a posterior distribution for *b*_*H*_. Details of each step are presented below. All steps are implemented in R (version 3.6.1) [14].

**Figure 2:**
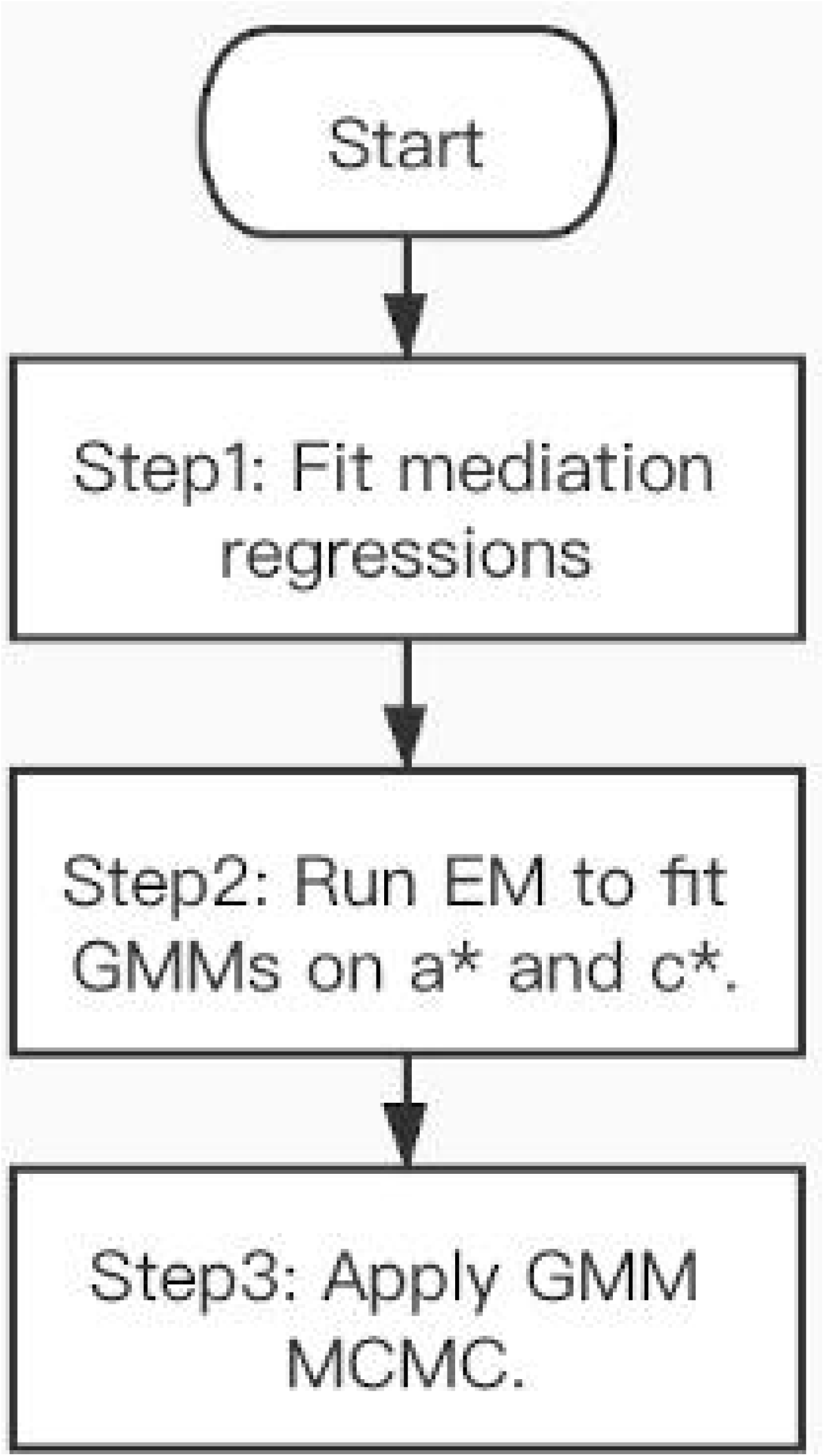
Flowchart of the multi-step method. In the first step, we use linear regression models to estimate the SNP effects on the known mediators (*a*^*^) and the direct effects between the SNPs and the outcome (*c*^*^). In the second step, we apply the EM algorithm to fit GMMs on ***a***^*^ and ***c***^*^. In the third step, a MCMC procedure was performed using the estimated GMM parameters from the last step in the priors to generate a posterior distribution for the hidden mediator’s effect size, *b*_*H*_.

#### 2.3.1 Step 1: Mediation regressions

In Step 1, we estimate the SNP effects on the known mediators, ***a***_**1**_ and ***a***_**2**_, and direct effects between the SNPs by fitting linear regressions for each known mediator separately where the mediator (e.g., *M*_1_) is the dependent variable and the elements of ***G*** are the independent variables. To estimate the direct effects between the SNPs and the outcome trait, ***c***, we fit a linear regression with *Y* being the dependent variable and ***G***, *M*_1_, *M*_2_, and other covariates being the independent variables. The resulting estimated effects are ***a***^⍰^ and ***c***^⍰^. We denote ***a***^⍰^ as the joint vector of ***a***^⍰^_**1**_ and ***a***^⍰^_**2**_.

#### 2.3.2 Step 2: EM

In Step 2, we separate the true effects in ***c***^⍰^ from the zero-mean noise component by fitting a GMM. In addition, because it is possible that not all the SNPs are associated with the set of known mediators, we also fit a GMM on ***a***^⍰^ to capture the actual effects of the SNPs on the known mediators. Specifically, we use the Expectation Maximization (EM) algorithm to fit the GMMs on ***a***^⍰^ and ***c***^⍰^ via the *normalmixEM* function in the *R* package *mixtools* (version 1.2.0) [7, 8, 9] with the initial value of mixing proportions, lambda, set to 0.5, which represents that initially, the SNPs have equal probabilities of being associated with the trait or not (i.e., the hidden mediator in the case of ***c***^⍰^ or the corresponding known mediators in the case of ***a***^⍰^).

The EM algorithm works as follows. Let *X*_*i*_ for *i* = 1…*n* be random variables generated from a GMM consist of two normal distributions (Equation 1), and let *Z*_*i*_ = {1,2} for *i* = 1…*n* be binary latent variables that each indicates which of the two normal distribution the corresponding *X*_*i*_ comes from.

Let 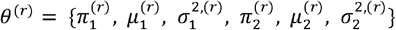 be the set of GMM parameters computed at iteration *r*. At iteration *r*, during the E step, compute *E*(*Z*_*i*_(*k*)|*x*_*i*_, θ^(*r*−1)^) = *P*(*Z*_*i*_ = *k*|*x*_*i*_, θ^(*r*−1)^), where *k* = {1,2} as in Equation 2. During the M step, compute elements of *θ*^(*r*)^as in Equations 3-5.

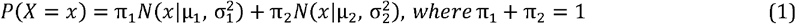

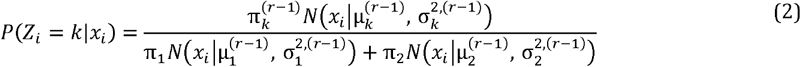

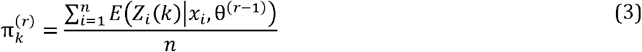

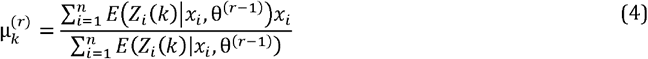

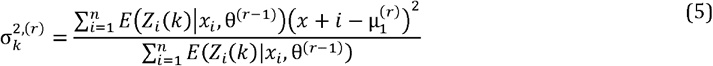

The fitted GMM should consist of a near zero-mean normal distribution, which are due to the zero-mean noise and a non-zero-mean normal distribution, which are due to the true effects. The means and the standard deviations of ***a***^⍰^’s and ***c***^⍰^’s true effect component can be estimated by the means and the standard deviations of the non-zero-mean distributions from the fitted GMMs. To avoid occasional convergence issues and extreme estimates, we run the EM algorithm multiple times and take the median value of the estimated true effect distribution means from all the runs. Based on our experience, generating 15 runs of the EM GMM is typically sufficient to obtain a well-fitted model.

#### 2.3.3 Step 3: GMM

In Step 3, we specify a GMM as shown in Equation 6 and perform a MCMC procedure to generate a posterior distribution for *b*_*H*_ [12, 13] using the R package *rjags* (version 4.10).

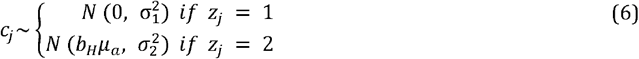

The data input into the GMM MCMC procedure are the elements of ***c***^⍰^. Denote the elements of the elements of ***c***^⍰^ as *c*_*j*_ for *j* = 1…*n*, where *n* is the number of SNPs in the model. 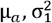, and 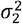 are constants in the model. The of *μ*_*a*_ is set to the estimated mean of the true effect Gaussian distribution of the fitted GMM on ***a***^⍰^ in Step 2; the value of 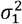 is set to the estimated variance of the zero-mean Gaussian distribution in the fitted GMM in step 2; the value of 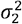 is set to the estimated variance of the non-zero-mean Gaussian distribution in the fitted GMM in step 2. The parameters estimated by this MCMC procedure are *b*_*H*_ and the binary indicator variables *z*_*j*_ for *j* = 1…*n*, which indicate which Gaussian distribution in the GMM the corresponding *c*_*j*_ belongs to. We specify the same categorical distribution prior for *z*_*j*_, where the weight parameters of the two categories are set to the corresponding estimated weights for the two Gaussian distributions in the fitted GMM in Step 2. Lastly, we specify a normal prior with mean zero and variance 100 for *b*_*H*_ so its distribution is almost flat at small values close to 0, which is the potential region of the hidden mediator’s effect size.

When *b*_*H*_ is equal to zero, the distribution for ***c***^⍰^ reduce to a single Gaussian distribution instead of a GMM, and the resulting GMM from EM will likely assign a very small weight to one of the Gaussian distributions in the GMM. If the Gaussian distribution that involves *b*_*H*_ in the MCMC procedure happens to be the one that receives a very small weight, the posterior distribution of *b*_*H*_ will span a very wide region around the true value of *b*_*H*_. This is because during each MCMC iteration, due to the small weight, very few or none of the elements in ***c***^⍰^ will be assigned to the Gaussian distribution that involves *b*_*H*_ so that the MCMC procedure is uncertain about the estimation of *b*_*H*_. This situation can also occur when the true value of *b*_*H*_ is very small such that the two Gaussian distributions in the GMM are not separable. Under these scenarios, the resulting interval estimators will be extremely wide and will not be useful in terms of giving a precise estimate of *b*_*H*_. Thus, if one observes an extremely wide posterior distribution of *b*_*H*_, we propose to flip the binary labeling of the fitted GMM from Step 2 and perform Step 3 again. Based on the results of the simulation study presented below, we observe that this adjustment generally results in more meaningful estimates of *b*_*H*_.

### 2.4 A simulation study

We conduct a simulation study to evaluate the proposed method. We consider a base case and seven additional settings. The base case is an ideal setting for our method. For each of the seven settings, we vary different aspects of the base case and evaluate the behavior of our method. In Setting 1, we vary *π*_*H*_, the frequency that the hidden mediator is associated with the exposure SNPs; in Setting 2, we allow two of the known mediators have negative effects on the outcome trait; in Setting 3, we decrease the sample size (number of individuals); in Setting 4, we consider cases where there are 1 and 10 known mediators of the outcome trait; in Setting 5, we consider four cases where some of the known mediators affect other known mediators; in Setting 6, we simulate two cases under the three-level SNP effect structure; in Setting 7, we simulate the case where *b*_*H*_ is equal to zero (i.e., negative control). For the base case and each of the seven settings, we consider a scenario where there are 70 SNPs associated with the outcome trait and another scenario where there are 500 SNPs associated with the outcome trait. For both scenarios, we let the hidden mediator’s effect size on the outcome trait, *b*_*H*_, be 0.02, 0.25, and 0.5 and perform 1000 independent simulations for each of the three values of *b*_*H*_. Also, for Settings 1-6, we simulate data for 49 additional *b*_*H*_ between 0.02 and 0.5 with a step size of 0.01 to show a behavior trend of our posterior distribution as *b*_*H*_ increases. As previously mentioned, when the true value of *b*_*H*_ is very small or equal to zero, the MCMC posterior distribution of *b*_*H*_ may be too wide occasionally to make any meaningful inference about its true value. Therefore, we apply the labeling-switching adjustment if the width of the 90% quantile interval derived from the initial posterior distribution for *b*_*H*_ is wider than 5. The detailed setting of the base case is presented in the next paragraph with its simulation results being presented in Section 3. The detailed settings and the simulation results of the other 7 settings are presented in the Supplementary Materials [see Additional file 1]. All simulations are performed in *R* (version 3.6.1) [14].

The data for the base case are simulated as follows. The sample size is 100,000. The SNPs associated with the outcome trait, ***G***, are simulated independently with minor allele frequencies generated from a uniform distribution between 0.1 and 0.5, and we assume the SNP effects are additive. The SNP effects on the mediators ***a***_**1**_, ***a***_**2**_, ***a***_**3**_, ***a***_**4**_, ***a***_**5**_ and ***a***_***H***_ are generated from mixture distributions of zero point-masses and normal distributions with the mean being 0.2 and the standard deviation being 0.08 as shown in Equations 7-12. The frequencies that the 5 known mediators are associated with the exposure SNPs are (0.5, 0.6, 0.8, 0.2, 0.5) (e.g., 50% of the SNPs associated with the outcome trait are associated with *M*_1_; 60% of the SNPs associated with the outcome trait are associated with *M*_2_, etc.). There are five known mediators of the outcome trait. The effect sizes of the five known mediators (*M*_1_, *M*_2_, *M*_3_, *M*_4_, *M*_5_) on the outcome trait are (0.4, 0.2, 0.3, 0.2, 0.4). The known mediators are generated as Equations 13-17. The frequency that the hidden mediator is associated with the exposure SNPs, *π*_*H*_, is 0.8. The hidden mediator is generated as Equation 18. We also include two covariates, *C*_1_ and *C*_2_, in the outcome model. *C*_1_ is generated from a normal distribution with a mean of 7 and a standard deviation of 0.5; *C*_2_ is generated from a normal distribution with a mean of 4 and a standard deviation of 0.4. Their corresponding effect sizes on the outcome trait are 0.8 and -0.3. The outcome trait is generated based on Equation 19. Note that we only add a relatively small error term (*ε*) to the outcome trait because we assume that almost all the leftover genetic heritability of the outcome trait can be explained by the hidden mediator. The mediators in the base case do not affect each other (i.e., no correlation among *M*_1_, *M*_2_, *M*_3_, *M*_4_, and *M*_5_).

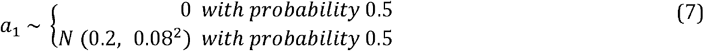

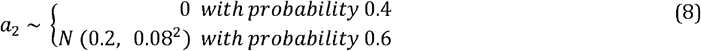

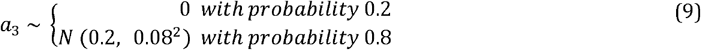

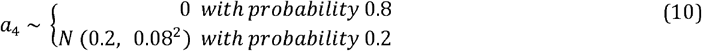

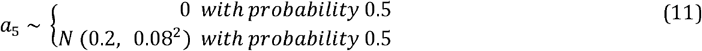

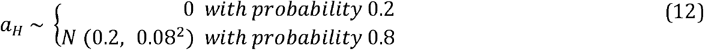

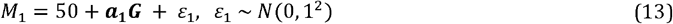

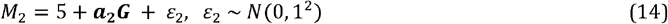

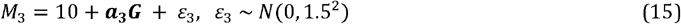

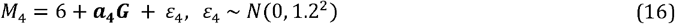

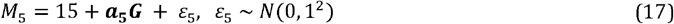

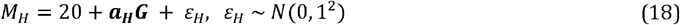

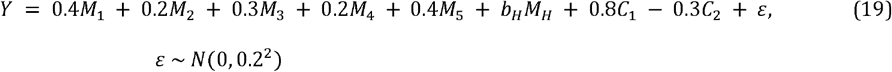

### 2.5 Data application

We apply the proposed method to UK Biobank data. We consider Waist-to-hip Ratio (WHR) as the outcome, the significant SNPs from the latest GWAS meta-analysis of WHR as the exposure, and body mass index (BMI) as the known causal mediator between WHR and the SNPs [15]. The goal of the analysis is to determine whether there exists a second, hidden mediator on WHR and if so, estimate the effect size of this hidden mediator. In addition to waist circumference, hip circumference and BMI, we also include sex, age, and the first ten genetic principal components in the UK Biobank phenotype data as covariates in the mediation model. Only individuals with European ancestry are considered for the current analysis, which includes those described as “British”, “Irish”, “White” or “Any other white background”. Individuals with missing phenotype data (i.e., “NA”) in any of the data fields are removed.

Because the GWAS meta-analysis used for identifying the exposure SNPs involves UK Biobank data, to minimize over-estimated SNP effect sizes in our analyses, we use a stringent p-value threshold of 5 × 10^−9^ when choosing the exposure SNPs. The resulting SNPs are clumped using the *R* package *TwoSampleMR* (version 0.5.6) with a clumping window of 250 kb and a cutoff for correlation due to linkage disequilibrium (LD *r*^2^ = 0.01) based on the 1000 Genomes Continental European groups reference [16, 19]. After LD clumping, a total of 535 independent SNPs associated with WHR are identified (Table S1) [see Additional file 2] and extracted from the imputed UK Biobank genotype using *plink* (version 2.0) [17, 18]. Furthermore, ten SNPs with low imputation quality (INFO score <0.9) are dropped from further analyses (labeled with “INFO” in Table S1), and a hard-call threshold of 0.4 is used when converting the imputed alleles probabilities to the number of allele copies. After joining the genotype data with the phenotype data, we further remove two SNPs with more than 10,000 missing rows (labeled with “NA” in Table S1) and a multi-allelic SNP (labeled with “M” in Table S1). Individuals with missing data (i.e., “NA”) in any of the data SNP fields are removed. In the end, 218,277 individuals and 522 SNPs are included in the application of our method. WHR is calculated as the ratio of waist circumference to hip circumference, and WHR and BMI are standardized to have means of zero and standard deviations of one.

The cleaned data are analyzed using the proposed multi-step method according to the flowchart in Figure 2. In Step 1, an initial regression was performed to access the effect direction of the SNPs on the outcome trait WHR using WHR as the dependent variable and all the SNPs, sex, age, and the first 10 genetic principal components as the independent variables. Based on the direction (positive or negative) of the estimated effect, the coding of each SNP is flipped such that all SNPs have a positive effect on WHR. In the subsequent mediator (BMI) regression models (to obtain ***a***^⍰^) and the outcome (WHR) regression model (to obtain ***c***^⍰^), sex, age and the first ten principal components are adjusted for as covariates. For both ***a***^⍰^ and ***c***^⍰^, SNPs with effects that are greater than the 3rd quartile + 3 × interquartile range (IQR) and values that are smaller than the 1st quartile - 3 × IQR are removed to avoid the downstream GMM methods to be driven by these outliers. A total of 12 SNPs were removed (label with “O” in Table S1). In Step 3, the MCMC chain length is set to 30,000 with a burn-in length of 5,000. We also estimate BMI’s effect on WHR by fitting a regression model with the dependent variable being WHR and the independent variables being BMI, sex, age and the first ten principal components.

## 3 Results

### 3.1 Simulation results

For the base case, simulation results on the median, mean, 90% highest density interval (HDI) and 90% quantile interval (QI) of *b*_*H*_’s posterior distributions are summarized in Figure 3. Rather than the usual 95% intervals, 90% intervals are reported to avoid the potential instability at the tails of the MCMC posterior distributions. We also report the root mean square error, the average bias, the number of outliers of the mean and the median point estimators, the proportion of the times that the HDI and QI contain the true value of *b*_*H*_, and empirical power / type I error of the HDI and the QI interval estimators for each of the eight simulation settings (Table S2, S3) [see Additional files 3 and 4]. The outliers are defined as the values more extreme than the third quartile + 1.5 * (the third quartile – the first quartile) or the first quartile – 1.5 * (the third quartile – the first quartile). For the base case and Settings 1-6, the empirical power is calculated as the number of simulations with the interval not containing zero divided by the total number of simulations. For Setting 7, the empirical type I error is calculated as the number of simulations with the interval not containing zero divided by the number of total simulations.

**Figure 3:**
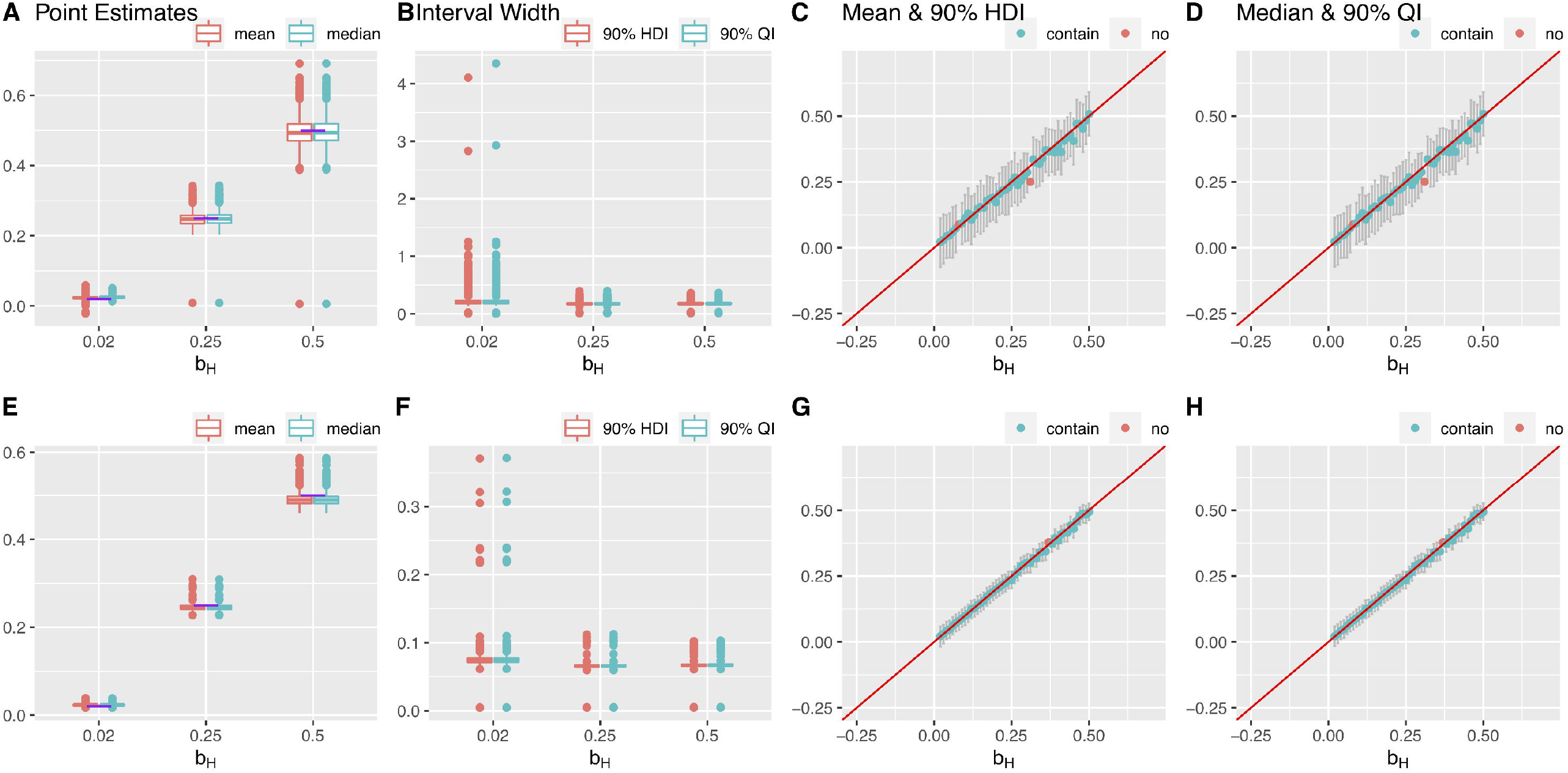
Results of one simulation setting: base case, varying *b*_*H*_, 1000 simulations. The first and second row presents the results for the 70 SNPs and 500 SNPs, respectively. (A, E) Box plots of the posterior median and the mean of *b*_*H*_. The purple lines indicate the true values. (B, F) Box plots of the widths of 90% HDIs and QIs. (C, G) The posterior medians and the 90% HDIs of the 49 equally spaced values of *b*_*H*_ between 0.02 and 0.5. (D, H) The posterior means and the 90% QIs of the 49 equally spaced values of *b*_*H*_ between 0.02 and 0.5. Outliers are defined as the values more extreme than the third quartile + 1.5 * (the third quartile – the first quartile) or the first quartile − 1.5 * (the third quartile – the first quartile).

Under the base case, the performances of the posterior median and mean of *b*_*H*_ are similar; both are close to the true value of *b*_*H*_, and slightly downward biased only when the true value of *b*_*H*_ is 0.5. Both the median and the mean have a smaller variation for the cases with 500 SNPs compared to 70 SNPs (Figure 3ACDEGH). In the case with 70 SNPs, for both *b*_*H*_ = 0.25 and *b*_*H*_ = 0.5, one out of the one thousand simulation runs, the median and the mean are far away from the corresponding true value (at values close to zero) (Figure 3A). But this is not observed in the 500 SNPs case. The 90% HDI and the 90% QI behave similarly as well; the intervals are wider for the cases with 70 SNPs than 500 SNPs only a few times, the HDI and the QI are extremely wide. For *b*_*H*_ = 0.02, *b*_*H*_ = 0.25 and *b*_*H*_ = 0.5 98.1%, 98.3%, and 97.3% of the 90% HDI and the 90% QI contains the true value of *b*_*H*_, indicating both intervals are conservative. In addition, when the true value of *b*_*H*_ is small, both interval estimators are slightly wider. Although the behavior of the posterior distribution of *b*_*H*_ does not change dramatically with the number of SNPs in the model, having more SNPs in the model can lead to slightly better estimations of *b*_*H*_ in terms of both the point estimators and the interval estimators.

Simulations were also conducted for seven additional settings. In Setting 1, where a low proportion of the SNPs are associated with the hidden mediator, the posterior median and mean shown slightly larger downward biases, and the HDI and the QI are wider when there is a smaller number of SNPs in the model. In contrast, when all of the SNPs are associated with the hidden mediators, the posterior median and mean are upward biased and the HDI and QI are less likely to capture the true value of *b*_*H*_ when the true value of *b*_*H*_ is large (Supplementary Materials Section 3.1) [see Additional file 1]. For Setting 2 and 3 such that the known mediators with negative effects on the outcome traits, simulation results show that varying the sample size (number of individuals) and the number of known mediators does not have dramatic impacts on the posterior distribution of *b*_*H*_ (Supplementary Materials Section 3.2, 3.3, 3.4) [see Additional file 1]. We also observe that in Setting 5, the posterior median and mean can be biased and the HDI and the QI are less likely to include the true value of *b*_*H*_ if the causal relationships among the known mediators are not appropriately adjusted for in the mediation regressions during Step 1 (Supplementary Materials Section 3.5) [see Additional file 1]. Furthermore, if the assumption that the SNP effects on all the mediators come from the same distribution does not hold, depending on the degree, the posterior median and mean can vary greatly and the HDI and the QI can have a low chance to include the true value of *b*_*H*_ as shown in Setting 6 (Supplementary Materials Section 3.6) [see Additional file 1]. Finally, according to the simulation result for Setting 7, the posterior median and mean are close to the true value of *b*_*H*_, zero, and the HDI and the QI have Type I error rates that are close to 0.1 (Supplementary Materials Section 3.7, Table S3) [see Additional files 1 and 4]. The detailed results of the additional settings are presented in the Supplementary Materials.

### 3.2 Application on waist-hip ratio

The regression estimated SNP effects on BMI (***a***^⍰^) and direct effects between the SNPs and WHR ***c***^⍰^ are shown in Figure 4.1A and C, respectively. As shown in the histograms, there are some extreme values or outliers in both ***a***^⍰^ and ***c***^⍰^. The histograms with outliers removed are shown in Figures 4.1B and 4.1D. Two modes can be observed from the distribution of ***a***^⍰^ with the mode on the left being approximately at zero suggesting ***a***^⍰^ follow a mixture model. It is less clear whether the distribution of ***c***^⍰^ has two modes. EM estimated means of ***a***^⍰^ and ***c***^⍰^ are indicated by the vertical red lines shown in Figure 4.1B and D. The fitted GMM on ***a***^⍰^ estimates a mixture weight of 0.882 for the distribution on the left, and the fitted GMM on ***c***^⍰^ estimates a mixture weight of 0.836 to the distribution on the left. From the posterior distribution of *b*_*H*_, the point estimate for *b*_*H*_ using the posterior median is 1.5556 and 1.5562 using the posterior mean; the HDI is (1.4658, 1.6440); the QI is (1.4683, 1.6469). The effect of BMI on WHR conditional on sex, age and the first ten principal components is estimated to be 0.3834 (95% CI: 0.3807, 0.3861).

**Figure 4:**
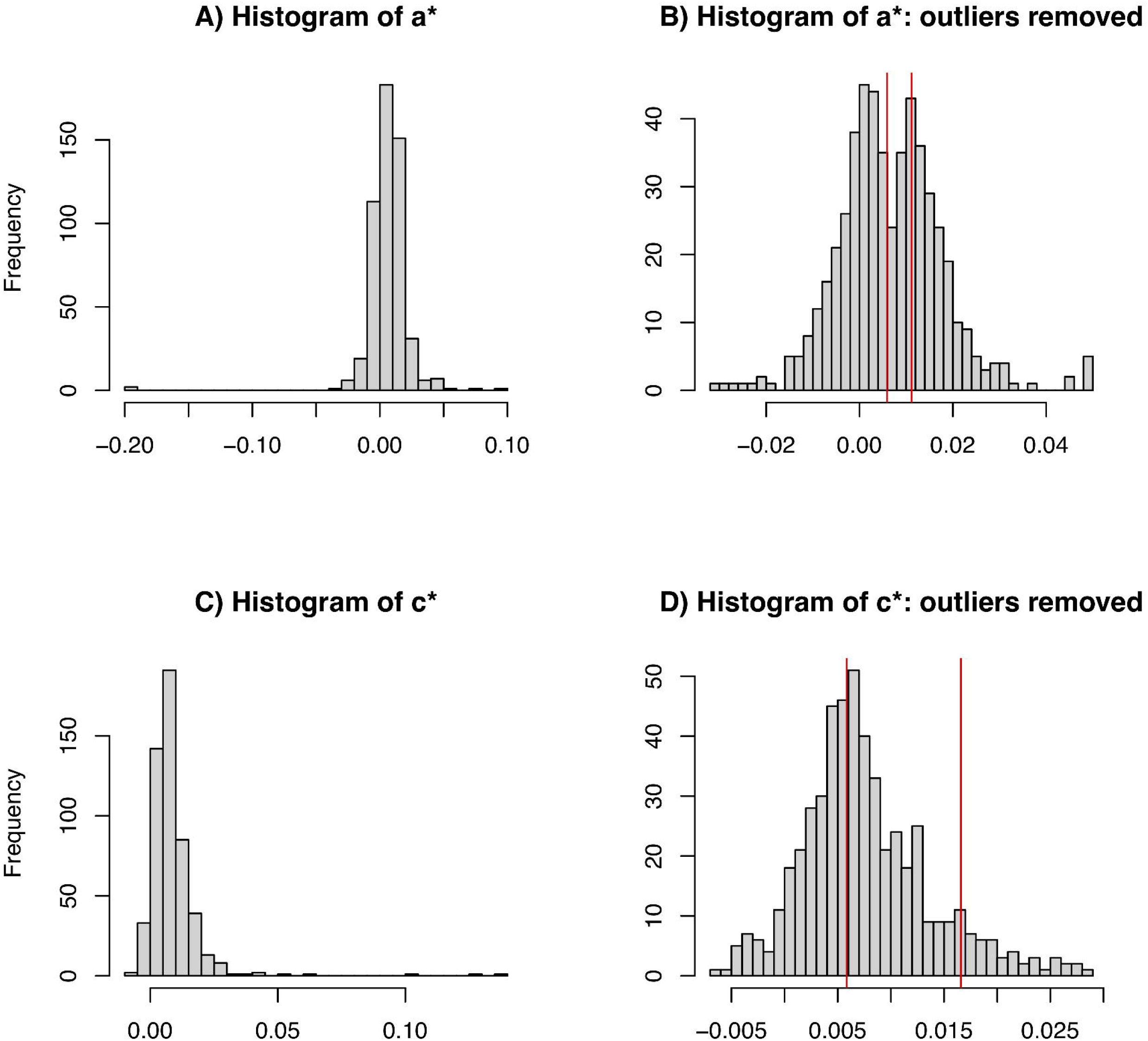
Histograms of the regression estimated effects. (A) Estimated SNP effects on BMI (*a*^⍰^). (B) Estimated SNP effects on BMI with outliers removed. (C) Estimated direct effects between the SNPs and WHR (*c*^⍰^). (D) Estimated direct effects between the SNPs and WHR with outliers removed.

## 4 Discussion

In this work, we propose to infer the effect size of a potential hidden mediator on a trait of interest based on the observed associations between the trait of interest, its known causal factors, and the associated SNPs that have been identified previously. Utilizing the mediation framework, we propose a multi-step method to estimate the effect size of the hidden factor by treating the trait of interest, its associated SNPs, and the known causal factors as the outcome, exposure, and known mediators in the mediation model, respectively. Assuming the direct effects between the outcome trait and the identified SNPs that are unexplained by the known mediators can be explained by a hidden mediator, we obtain the effect of this hidden mediator on the outcome trait by decomposing the direct effects between the outcome trait and its associated SNPs into the SNP effects on the hidden mediator and the hidden mediator’s effect size on the outcome trait. In Step 1 of our proposed method, we estimate the SNP effects on the known mediators and the direct effects between the SNPs and the outcome trait via a series of linear regressions. In Step 2, we fit two GMMs on the estimated SNP effects on the known mediators and the direct effects between the SNPs and the outcome trait using EM algorithm to separate the actual SNP effects from the zero-mean noises (i.e., the estimated effects of those SNPs not associated with the known and hidden mediators). Lastly in Step 3, based on the EM estimated GMM parameters, a GMM MCMC procedure is applied to generate a posterior distribution for the hidden mediator’s effect size, *b*_*H*_.

Through extensive simulation studies, we show that our method can produce a posterior distribution that captures *b*_*H*_ well. We observe that if the model assumptions are correct, both using the posterior median and mean to estimate *b*_*H*_ provides only small biases, and good coverages by the 90% HDI and QI. When there are more identified SNPs associated with the outcome trait, in general, both the point and the interval estimators perform better. Also, our posterior distribution estimates the hidden mediator’s effect size well if some of the known mediators have negative effects on the outcome trait while others have positive effects. Our method can also accommodate varying numbers of known mediators and the performance is not dramatically affected by decreasing sample size. In addition, our method estimates *b*_*H*_ well even if the hidden mediator does not exist (i.e., *b*_*H*_ =0). However, we notice that when the hidden mediator is associated with all of the identified SNPs that are associated with the outcome trait, both the median and the mean are upward biased and both the HDI and the QI are less likely to include the true value of *b*_*H*_. This is expected because the assumed GMM distribution on the estimated direct effects between the SNPs and the outcome trait is wrong. On the other hand, if the hidden mediator is associated with too few identified SNPs, the point estimators have large biases and the interval estimators become wider. This is also expected as there are fewer SNPs that can provide information about *b*_*H*_. Also, if causal relationships exist among the known mediators, these effects need to be adjusted in the mediation regressions. Otherwise, the point estimators can have relatively large biases and the interval estimators are likely to include the true value if the causal relationships are large enough. Finally, our method assumes that the SNP effects on different mediators (including both the known and hidden mediators) are similar. The posterior distribution estimates *b*_*H*_ poorly when the SNP effects on different mediators are very different, especially when the number of SNPs in the model is large. This is because little information can be borrowed from the observed SNP effects to infer the SNP effects on the hidden mediator, which can make it difficult to decompose the direct effects between the outcome trait and the associated SNPs to estimate *b*_*H*_.

Occasionally, a posterior distribution with little information about *b*_*H*_ (i.e., is extremely wide) can be generated when the true value of *b*_*H*_ is either zero or very close to zero such that the distribution of the estimated direct effects between the outcome trait and the associated SNPs follows a single Gaussian distribution around zero rather than a GMM. Under this scenario, by chance alone, the EM algorithm in Step 2 may assign a tiny weight to the distribution that involves *b*_*H*_ in the MCMC model in Step 3 and assign a large weight to the other distribution in the GMM. As a result, little data can be used to infer *b*_*H*_ in the MCMC procedure. To avoid this situation, we suggest one inspect the histogram of estimated direct effects and the EM fitted GMM from Step 2. If the histogram of estimated direct effects does not have two modes and is centered approximately at zero, and the EM fitted GMM assigns a tiny weight to the distribution that involves *b*_*H*_ relative to the weight of the other distribution, then it is reasonable to flip the binary labeling of GMM and proceed to Step 3.

We applied our proposed method on UK Biobank data to estimate the effect size for a potential hidden mediator of waist-hip ratio. From the posterior distribution generated by our method, the posterior median estimates that a potential hidden mediator exists in the European population with an effect size of 1.56 (90% QI: 1.47, 1.64). This result suggests that the hidden mediator has a larger effect on waist-hip ratio comparing to BMI (0.38). Some caution here is warranted, as we used the same UK Biobank data for both identifying SNPs associated with waist-hip ratio and for estimating the SNP effects. Although we used an extra stringent p-value threshold (5 × 10^−9^) for filtering SNPs associated with waist-hip ratio to mitigate biases from winner’s curse, some degree of biases from the winner’s curse is unavoidable. A more optimal approach is to identify SNPs associated with waist-hip ratio and perform the estimation in two independnent populations with similar ancestries such that the effects are similar in the two populations but the biases from the winner’s curse are minimal.

Our proposed method has some limitations. First, the performance of the posterior distribution for *b*_*H*_ under our method largely depends on the how well we estimate the regression coefficients for the mediation regressions during Step 1, as the downstream steps treat the estimated regression coefficients as input data. Precise and accurate estimates of the coefficients require the mediation regression to be performed on data sets with large samples size, especially when many SNPs are included in the model. For large population genetics data, this may be less of a concern. As we learned from the simulation studies, the posterior distribution captures *b*_*H*_ well when the sample size is 25,000, which for today’s conventional size of DNA biobanks is not unreasonable. However, if strong correlations among the known mediators are not properly adjusted in the mediation regressions, the regression coefficients will be biased, which can lead to substantial biases in the resulting posterior of *b*_*H*_. Thus, our method relies on one’s input of prior domain knowledge and the specification of reasonable regression models. Future work can be devoted to extending the methods to address the situation when the known mediators are correlated with each other and no prior knowledge on the causal directions among the known mediators is available. Second, our method relies on the strong assumption that the SNP effects on all the mediators between the SNPs and the outcome trait come from the same distribution, and departure from this assumption can lead to substantial variation for the posterior distribution for *b*_*H*_ such that the inference based on the posterior distribution can be very inaccurate. However, we argue that it is somewhat reasonable to constrain the SNP effects on the hidden mediator to be similar to the SNP effects on the known mediators and there will be infinite number of ways to decompose the direct effect between the SNPs and the outcome trait without this assumption. Future work can focus on relaxing this assumption or inventing ways to incorporate information about the potential SNP effect sizes from other modality. Furthermore, as an initial step to learn about what we do not know about an outcome trait, we assume that all the leftover associations between the outcome trait and its association SNPs are due to one hidden mediator. In reality, any leftover associations – and perhaps more likely – could be due to multiple hidden mediators each with a relatively small effect size. Even if there are multiple hidden mediators, we can interpret the estimated effect size assuming a single hidden mediator as the combined effect of multiple potential hidden mediators. Future work should develop a method to infer the number of potential hidden mediators and decompose the combined effect into individual effects from each of the hidden mediators. Lastly, future work can extend the continuous outcome trait to other outcome types such as binary variables potentially via the counterfactual framework. This will involve utilizing generalized linear models in the meditation regressions in Step 1. Such extension will make the method useful for many applications such that the disease trait is binary.

## 5 Conclusions

We developed a method for estimating the effect size of a potential hidden estimator between A trait of interest and its associated SNPs. In the first step, a series of regression models are used to estimate the SNP effects on the hidden mediators and the direct effects between the SNPs and the trait of interest. In the second step, GMM models are fitted to the estimated SNP effects on the hidden mediators and the estimated direct effects between the SNPs via the EM algorithm. In the final step, an MCMC procedure that utilizes parameters estimated in the second step is used for generating a posterior distribution for the hidden mediator’s effect size. Extensive simulations show that our method can generate accurate estimators for the hidden mediator’s effect size. Also, when the hidden mediator does not exist, our method has controlled type I error rates. By applying our method to UK Biobank data, we found a potential hidden mediator between waist-hip-ratio and its associated SNPs in the European population and estimated its effect size on waist-hip-ratio to be larger than a known mediator BMI’s effect size on waist-hip-ratio. Although, as an initial step toward finding the hidden mediators between a trait of interest and its associated SNPs, we hypothesize a simple model with only one hidden mediator left, we hope that our method can provide some insights into the characteristics of the remaining one or multiple hidden mediators and inspire further method developments in this direction.

## Supporting information

Additiona_file_1

Additiona_file_2

Additiona_file_3

Additiona_file_4

## 5 Declarations

### 5.1 Ethics approval and consent to participate

We confirm that all methods were carried out in accordance with relevant guidelines and regulations.

### 5.2 Consent for publication

Not applicable.

### 5.3 Availability of data and materials

The UK Biobank data are provided under application ID 32133 and can be accessed by others at the UK Biobank website (https://www.ukbiobank.ac.uk). R Scripts with the simulation code and an R package that implements the described method are available at https://github.com/zhd007/HiddenMediator [19].

### 5.4 Competing interests

The authors have no conflicts to report.

### 5.5 Funding

B.F.V acknowledges support from the National Institutes of Health (DK101478, DK126194) and a Linda Pechenik Montague Investigator Award. MDR acknowledges support from the National Institutes of Health (AI077505).

### 5.6 Author’s contributions

Z.D. and B.F.V. conceived of the project and designed the experiments. M.D.R. provided data. Z.D. analyzed the data. Z.D., W-T.H., and B.F.V. developed the method and prepared a draft of the initial manuscript. All authors edited the manuscript. W-T.H and B.F.V. supervised the project.

## 5.7 Acknowledgements

Not applicable.

## 5.8 Authors’ information

Voight, B.F. and Hwang, W-T are joint corresponding authors.

## 8 Additional material information

File name: Additional_file_1.pdf

File format: .pdf

Title of data: Supplementary Materials

Description of data: A pdf file containing supplementary text (Sections S1-S4) and figures (Figures S1-S20) including the discussion on a more generation assumption on the SNP effects (Section S1), additional simulation settings and results (Sections S2-S3), and a proof of the estimated SNP effect’s distribution (Section S4).

File name: Additional_file_2.xlsx

File format: .xlsx

Title of data: Table S1, a list of independent SNPs associated with WHR

Description: An Excel table containing the list of 535 independent SNPs associated with WHR.

File name: Additional_file_3.xlsx

File format: .xlsx

Title of data: Table S2, simulation results for Settings 1-6

Description: An Excel table containing the simulation results for Settings 1-6.

File name: Additional_file_4.xlsx

File format: .xlsx

Title of data: Table S3, simulation result for Setting 7

Description: An Excel table containing the simulation results for Setting 7.

## 9 Contact information

Zhuoran Ding

Marylyn D. Ritchie

Benjamin F. Voight

Wei-Ting Hwang

## Notes

### Competing Interest Statement

The authors have declared no competing interest.

