## Supplementary material for "Estimating the effect size of a hidden causal factor between SNPs and a continuous trait: a mediation model approach": Additiona_file_1

### Supplementary Materials

**S4 Proof: The distribution of $(a^*_{ij}|\Delta_{ij} = 1)$ is $N(\mu_u, \sigma^2_u + \sigma^2_a + \sigma^2_{a^*_{ij}})$.** 27

**S5 Figure legends** 29

### S1 A more general assumption on SNP effects

As previously mentioned, the assumption that different SNP effects on mediators come from the same distribution may be strong. In this section, we introduce a three-level structure for the SNP effects as an alternative framework. We will evaluate our proposed method by simulating data accordingly. Let $i$ be the index for mediators and $j$ be the index for SNPs. The three-level structure of the SNP effects on mediators $\boldsymbol{a}$ and the direct effects $\boldsymbol{c}$ is as follow:

For the SNP effects on known mediators, $\boldsymbol{a}$:

Level 1: $M_i$'s mediator-specific mean, $u_i$

$$u_i \sim N(\mu_u, \sigma_u^2) \tag{S1}$$

Level 2: the effect of SNP $j$ on the mediator $i$, $a_{ij}$

$$a_{ij}|u_i \sim \begin{cases} 0 & \text{with probability } 1-\pi_i \\ N(u_i, \sigma_a^2) & \text{with probability } \pi_i \end{cases} \tag{S2}$$

Level 3: the regression estimated effect of SNP $j$ on the mediator $i$, $a^*_{ij}$

$$a^*_{ij} \sim \begin{cases} N(0, \sigma^2_{a^*_1}) & \text{with probability } 1-\pi_i \\ N(\mu_u, \sigma^2_{a^*_2}) & \text{with probability } \pi_i \end{cases} \tag{S3}$$

For the direct effect SNPs on the outcome, $\boldsymbol{c}$:

Level 1: mediator-specific mean, $u_i$

$$u_i \sim N(\mu_u, \sigma_u^2) \tag{S4}$$

Level 2: the direct effect between SNP $j$ and the outcome trait, $c_j$

$$c_j|u_i \sim \begin{cases} 0 & \text{with probability } 1-\pi_H \\ N(b_H u_i, \sigma_c^2) & \text{with probability } \pi_H \end{cases} \tag{S5}$$

Level 3: the regression estimated direct effect between SNP $j$ and the outcome trait, $c^*_j$

$$c^*_j \sim \begin{cases} N(0, \sigma^2_{c^*_1}) & \text{with probability } 1-\pi_H \\ N(b_H \mu_u, \sigma^2_{c^*_2}) & \text{with probability } \pi_H \end{cases} \tag{S6}$$

Figure S1 illustrates the case when the first two levels for the effects of the SNPs that are associated with the mediator (excluding the non-effect SNPs) in a three-mediator ($M_1$, $M_2$, $M_H$) model. Figure S1B shows the second level in which SNP effects on each mediator, including the hidden mediator, are assumed to come from a mediator-specific normal distribution in colors of yellow, blue, and green. The means of the mediator-specific normal distributions are assumed to come from the same normal distribution in Level 1 as shown in the red curve in Figure S1A. Based on the three-level structure, the mean SNP effect of a specific mediator $M_i$ is $u_i$. This allows SNP effects on different mediators to be different. The differences among SNP effects on different mediators can be represented by the normal distribution at Level 1 with variance $\sigma_u^2$. If $\sigma_u^2$ is zero, then the model reduces to the original assumption that the SNP effects on different mediators are the same. Level 2 specifies the distribution of $\boldsymbol{a}$ and $\boldsymbol{c}$, which are mixture distributions of the effect sizes of SNPs associated with the mediator and the effect sizes of SNPs that have zero effect on the mediator. Level 3 specifies the distributions of regression estimated effects, $\boldsymbol{a}^*$ and $\boldsymbol{c}^*$, and the point mass at zero becomes a normal distribution due to the estimation uncertainty. Since the actual effect (non-zero centered) distribution of $\boldsymbol{a}^*$ is centered at $\mu_u$ and the actual effect (non-zero centered) distribution of $\boldsymbol{c}^*$ is centered at $b_H\mu_u$, we can estimate $\mu_u$ from $\boldsymbol{a}^*$ and $b_H\mu_u$ from $\boldsymbol{c}^*$. Consequently, estimates of $b_H$ can be obtained by the proposed MCMC procedure. The derivations of the Level 3 distributions are provided in the Supplementary Material Section 4.

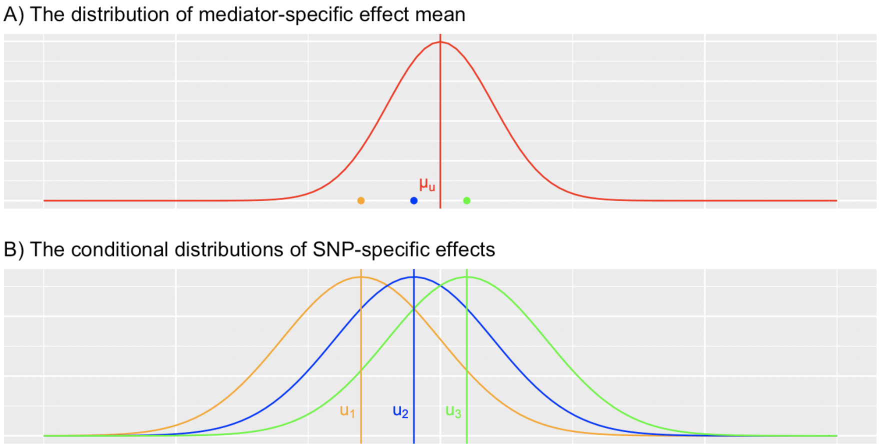

Figure S1: The first two levels of the three-level SNP effect structure. A) The distribution of mediator-specific effect mean; B) The conditional distribution of SNP-specific effects.

### S2 Additional simulation settings

We present the seven additional simulation settings in this section. Each of the seven settings differs from the base case presented in the main text in some aspect then we evaluate the impact of a particular aspect on the performance of the proposed method.

#### S2.1 Setting 1: different frequencies of the hidden mediator

In Setting 1, we set $\pi_H$, the frequency that the hidden mediator is associated with the exposure SNPs, to be 1, 0.5 and 0.3. In the base case, $\pi_H = 0.8$.

##### S2.2 Setting 2: negative effects

In Setting 2, we set two of the known mediator's effect sizes to be negative such that the known mediator's effect sizes are (0.4, 0.2, 0.3, -0.2, -0.4).

##### S2.3 Setting 3: different sample sizes

In Setting 3, we decrease the sample size to be 50,000 and 25,000. The sample size in the base case is 100,000.

##### S2.4 Setting 4: different numbers of mediators

In Setting 4, we vary the number of known mediators in the model. We consider two additional cases: one mediator and ten mediators. In both cases, the hidden mediator $M_H$, the SNP effects $\boldsymbol{a_H}$ and $M_H$'s effect on the outcome trait $b_H$ are generated in the same way as in the base case. In the one-mediator case, the known mediator $M_1$, SNP effect $\boldsymbol{a_1}$ and $M_1$'s effect on the outcome trait are generated in the same way as the first mediator $M_1$ in the base case. The outcome $Y$ is generated as in Equation S7. In the ten-mediator case, the first 5 mediators ($M_1$, $M_2$, $M_3$, $M_4$, $M_5$), the SNP effects on the first 5 mediators ($\boldsymbol{a_1}$, $\boldsymbol{a_2}$, $\boldsymbol{a_3}$, $\boldsymbol{a_4}$, $\boldsymbol{a_5}$), and their effects on the outcome trait are generated in the same way as those of the first 5 mediators in the base case. The SNP effects on the 5 additional mediators, $M_6$, $M_7$, $M_8$, $M_9$ and $M_{10}$, are generated in the same form as in Equations 2-6 in the main text, but the frequencies that $M_6$, $M_7$, $M_8$, $M_90$ and $M_{10}$ are associated with the exposure SNPs are 0.8, 0.3, 0.4, 0.5, 0.6, respectively. The values of $M_6$, $M_7$, $M_8$, $M_9$, $M_{10}$ and the outcome $Y$ are generated as Equations S8-S13.

$$Y = 0.4 + 0.8C_1 - 0.3C_1 + \epsilon,\, \epsilon \sim N(0,\, 0.2^2) \tag{S7}$$

$$M_6 = 15 + \boldsymbol{a_6 G} + \epsilon_6,\, \epsilon_6 \sim N(0,\, 1.5^2) \tag{S8}$$

$$M_7 = 10 + \boldsymbol{a_7 G} + \epsilon_7,\, \epsilon_7 \sim N(0,\, 1^2) \tag{S9}$$

$$M_8 = 5 + \boldsymbol{a_8 G} + \epsilon_8,\, \epsilon_8 \sim N(0,\, 0.5^2) \tag{S10}$$

$$M_9 = 2 + \boldsymbol{a_9 G} + \epsilon_9,\, \epsilon_9 \sim N(0,\, 0.5^2) \tag{S11}$$

$$M_{10} = 20 + \boldsymbol{a_1 0G} + \epsilon_{10},\, \epsilon_{10} \sim N(0,\, 1^2) \tag{S12}$$

$$
\begin{aligned}
Y = & \; 0.4M_1 + 0.2M_2 + 0.3M_3 + 0.2M_4 + 0.4M_5 \\
& + 0.2M_6 + 0.1M_7 + 0.3M_8 + 0.2M_9 + 0.2M_{10} \\
& + b_H M_H + 0.8C_1 - 0.3C_1 + \epsilon, \; \epsilon \sim N(0, \, 0.2^2)
\end{aligned} \tag{S13}
$$

#### S2.5 Setting 5: correlated mediators

In Setting 5, the causal relationships among the known mediators are allowed. We consider four cases. (1) $M_1$ affects $M_2$ with an effect size of 0.3. (2) $M_1$ affects $M_2$ with an effect size of 0.9. (3) $M_1$ affects $M_2$ with an effect size of 0.3, and $M_3$ affects $M_4$ with an effect size of 0.2. (4) $M_1$ affects $M_2$ with an effect size of 0.5, and $M_3$ affects $M_4$ with an effect size of 0.4. For all four cases, we first generate all the mediators in the same way as in the base case, then add the product of a mediator and its corresponding effect size to the mediator that it affects. For example, in the first case, we generate $M_1$, $M_2$, $M_3$, $M_4$, $M_5$ in the same way as in the base case, then add 0.3 times $M_1$ to $M_2$. We first ignore the causal relations among the known mediators in the mediation regressions during Step 1 of our method, apply the usual Step 2 and 3, and evaluate the results. Next, we evaluate the results where the causal relations among the known mediators in the mediation regressions during Step 1 are correctly adjusted for. For example, in the first case, we adjust for the association between $M_1$ and $M_2$ by including $M_1$ as an independent variable in the regression model where $M_2$ is the dependent variable. Step 2 and Step 3 remain the same as in the base case.

#### S2.6 Setting 6: the three-level SNP effect structure

In the base case, SNP effects on all six mediators are generated from a normal distribution with a mean of 0.2 and a standard deviation of 0.08. Under the three-level SNP effect structure, this is equivalent to setting the Level 1 standard deviation (the difference among the mediators) to 0 and the Level 2 standard deviation to 0.08. In Setting 6, we generate the SNP effects under the more general three-level SNP effect assumption mentioned in Supplementary Material Section 1. Specifically, we fix the sum of the Level 1 standard deviation (i.e., $\sigma_u$ in Equation S1) and the Level 2 standard deviation (i.e., $\sigma_a$ in Equation S2) to be 0.08 and vary the ratio of the Level 1 and Level 2 standard deviations. We let the Level 1 and Level 2 standard deviation ratios be 1:3 and 1:1. The corresponding Level 1 and Level 2 standard deviations are (0.02, 0.06) and (0.04, 0.04). Using the 1:3 ratio scenario as an example, we simulate the three-level effect structure by first generating six mediator-specific means for $M_1$, $M_2$, $M_3$, $M_4$, $M_5$ and $M_H$ from a normal distribution with a mean of 0.2 and a standard deviation 0.02, then we generate SNP effects on each mediator from a normal distribution with the mean being the mediator's mediator-specific mean and the standard deviation being 0.06.

##### S2.7 Setting 7: no hidden mediator

In Setting 7, we set $b_H$ to 0 and evaluate the proposed method when there is no hidden mediator between the outcome trait and the associated SNPs.

#### S3 Additional simulation results

##### S3.1 Setting 1: different frequencies of the hidden mediator

In Setting 1, we evaluate our method under different $\pi_H$, where $\pi_H$ is the frequency that the SNPs are associated with the hidden mediator. Specifically, we let $\pi_H$ equal 0.3, 0.5, and 1, whereas the base case $\pi_H$ is 0.8. First, when $\pi_H$ is less than 1, in the 70 SNP case, the HDI and the QI are wider when $\pi_H$ is smaller (Figures S2BCD, S3BCD). In the 500 SNP case, this trend is less obvious (Figures S2FGH, S3FGH). This is because a smaller $\pi_H$ leads to a smaller number of observations of the direct effects between the SNPs associated with the hidden mediator and the outcome trait. With a fewer number of direct effect estimates available for Step 3, the MCMC posterior distribution is less certain about the potential value of $b_H$. For example, if there are 70 SNPs associated with the outcome trait, and $\pi_H$ equals 0.3, it implies that there are only about 21 SNPs associated with the hidden mediator to be used for inferring $b_H$. On the other hand, when there are 500 SNPs, even if $\pi_H$ is 0.3, there are still 150 SNPs associated with the hidden mediator for making a confident estimation of $b_H$. Also, the mean and the median have more downward biases as $\pi_H$ decreases, especially when the true value of $b_H$ is 0.5 (Figures S2AE, S3AE, Table S1). When $\pi_H$ is equal to 1, the mean and the median are upward biased, and the HDI and the QI are less likely to contain the true values, especially for larger values of $b_H$, which shows that the posterior distribution overestimates $b_H$ (Figure S4, Table S1). This is because when $\pi_H$ is 1, rather than following the assumed GMM, $\boldsymbol{c}^*$ follow a single Gaussian distribution. But the EM algorithm in Step 2 will try to separate the $\boldsymbol{c}^*$ into two distributions and use the larger-value distribution of the two for inferring $b_H$ so the resulting posterior distribution will overestimate the true value of $b_H$.

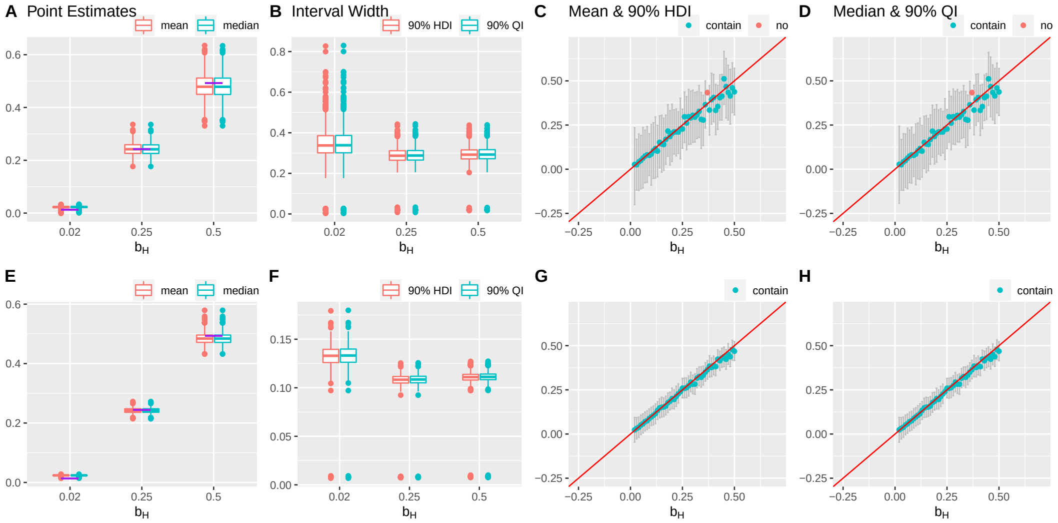

Figure S2: Results of a simulation setting: $\pi_H = 0.3$, varying $b_H$, 1000 simulations. The first and second row presents the results for the 70 SNPs and 500 SNPs, respectively. (A, E) Box plots of the posterior median and the mean of $b_H$. The purple lines indicate the true values. (B, F) Box plots of the widths of 90% HDIs and QIs. (C, G) The posterior medians and the 90% HDIs of the 49 equally spaced values of $b_H$ between 0.02 and 0.5. (D, H) The posterior means and the 90% QIs of the 49 equally spaced values of $b_H$ between 0.02 and 0.5. Outliers are defined as the values more extreme than the third quartile + 1.5 * (the third quartile – the first quartile) or the first quartile – 1.5 * (the third quartile – the first quartile).

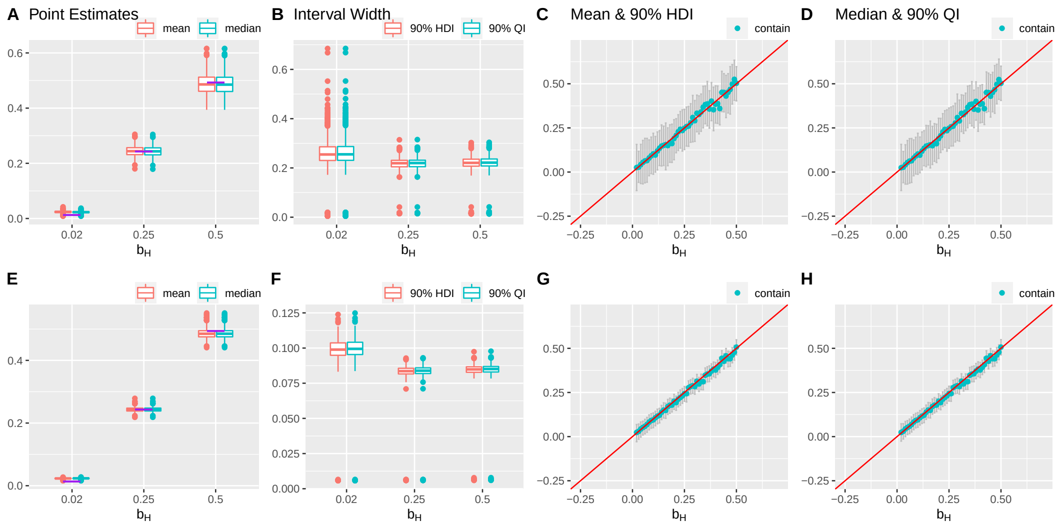

Figure S3: Results of a simulation setting: $\pi_H = 0.5$, varying $b_H$, 1000 simulations. The first and second row presents the results for the 70 SNPs and 500 SNPs, respectively. (A, E) Box plots of the posterior median and the mean of $b_H$. The purple lines indicate the true values. (B, F) Box plots of the widths of 90% HDIs and QIs. (C, G) The posterior medians and the 90% HDIs of the 49 equally spaced values of $b_H$ between 0.02 and 0.5. (D, H) The posterior means and the 90% QIs of the 49 equally spaced values of $b_H$ between 0.02 and 0.5. Outliers are defined as the values more extreme than the third quartile + 1.5 * (the third quartile – the first quartile) or the first quartile – 1.5 * (the third quartile – the first quartile).

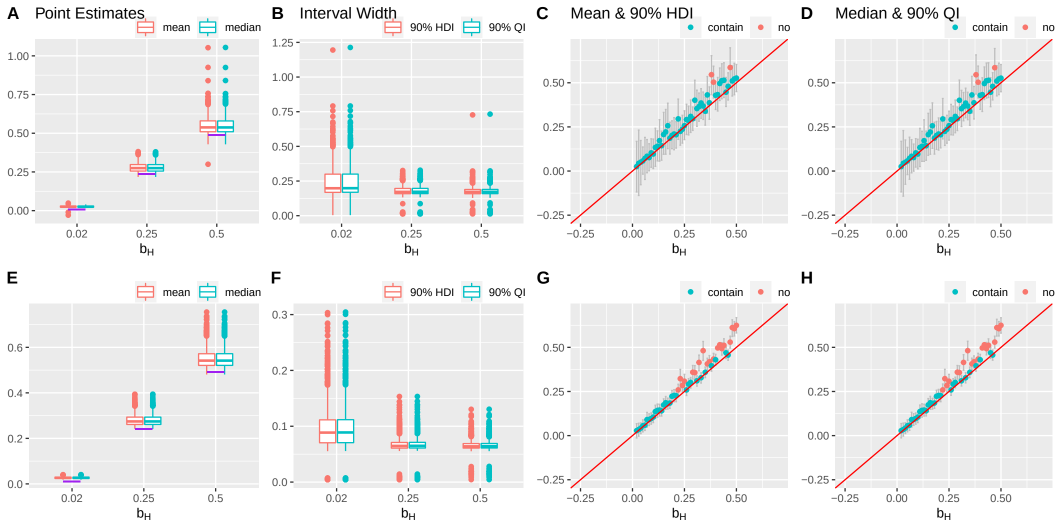

Figure S4: Results of a simulation setting: $\pi_H = 1$, varying $b_H$, 1000 simulations. The first and second row presents the results for the 70 SNPs and 500 SNPs, respectively. (A, E) Box plots of the posterior median and the mean of $b_H$. The purple lines indicate the true values. (B, F) Box plots of the widths of 90% HDIs and QIs. (C, G) The posterior medians and the 90% HDIs of the 49 equally spaced values of $b_H$ between 0.02 and 0.5. (D, H) The posterior means and the 90% QIs of the 49 equally spaced values of $b_H$ between 0.02 and 0.5. Outliers are defined as the values more extreme than the third quartile + 1.5 * (the third quartile – the first quartile) or the first quartile – 1.5 * (the third quartile – the first quartile).

#### S3.2 Setting 2: negative effects

In Setting 2, two out of five known mediators have negative effects on the outcome trait. Based on the simulation results, the behavior of both the point estimators and the interval estimators under this setting is similar to that of the base case (Figure S5, Table S1).

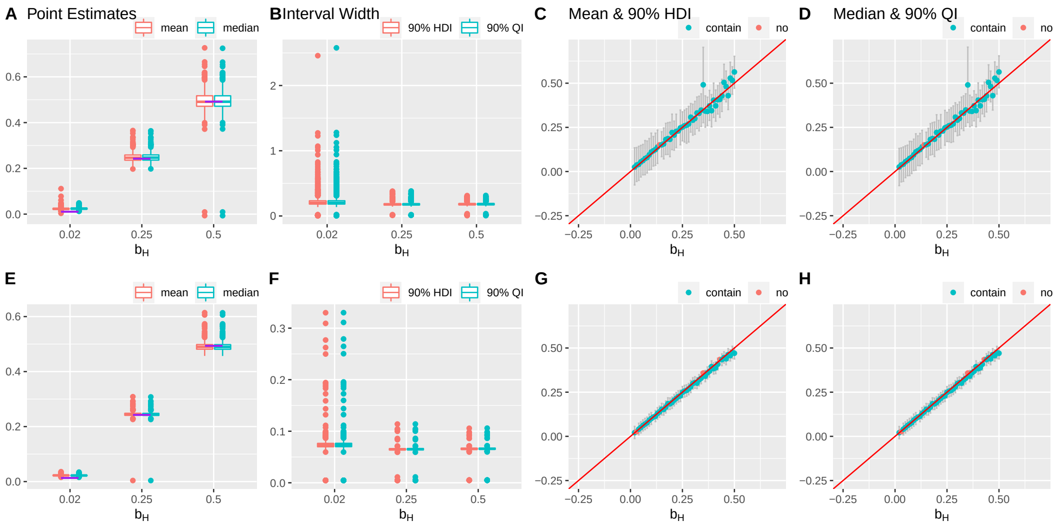

Figure S5: Results of a simulation setting: negative effects, varying $b_H$, 1000 simulations. The first and second row presents the results for the 70 SNPs and 500 SNPs, respectively. (A, E) Box plots of the posterior median and the mean of $b_H$. The purple lines indicate the true values. (B, F) Box plots of the widths of 90% HDIs and QIs. (C, G) The posterior medians and the 90% HDIs of the 49 equally spaced values of $b_H$ between 0.02 and 0.5. (D, H) The posterior means and the 90% QIs of the 49 equally spaced values of $b_H$ between 0.02 and 0.5. Outliers are defined as the values more extreme than the third quartile + 1.5 * (the third quartile – the first quartile) or the first quartile – 1.5 * (the third quartile – the first quartile).

#### S3.3 Setting 3: different sample sizes

In Setting 3, we evaluate our method when the sample size (i.e., the number of individuals in the study) is smaller. Specifically, we reduce the sample size to 50,000 and 25,000, whereas the base case sample size is 100,000. With smaller sample sizes, both the point estimators and the interval estimators behave similarly to the base case (Figure S6, S7, Table S1). Only the HDI and the QI are slightly wider when the sample size is smaller (Figure S6BF, S7BF).

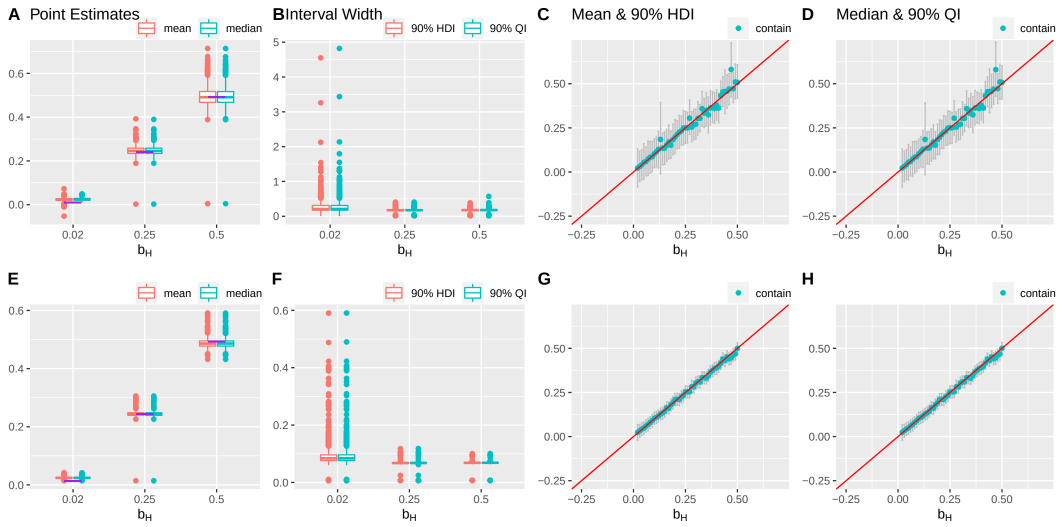

Figure S6: Results of a simulation setting: $n$ = 50,000, varying $b_H$, 1000 simulations. The first and second row presents the results for the 70 SNPs and 500 SNPs, respectively. (A, E) Box plots of the posterior median and the mean of $b_H$. The purple lines indicate the true values. (B, F) Box plots of the widths of 90% HDIs and QIs. (C, G) The posterior medians and the 90% HDIs of the 49 equally spaced values of $b_H$ between 0.02 and 0.5. (D, H) The posterior means and the 90% QIs of the 49 equally spaced values of $b_H$ between 0.02 and 0.5. Outliers are defined as the values more extreme than the third quartile + 1.5 * (the third quartile – the first quartile) or the first quartile – 1.5 * (the third quartile – the first quartile).

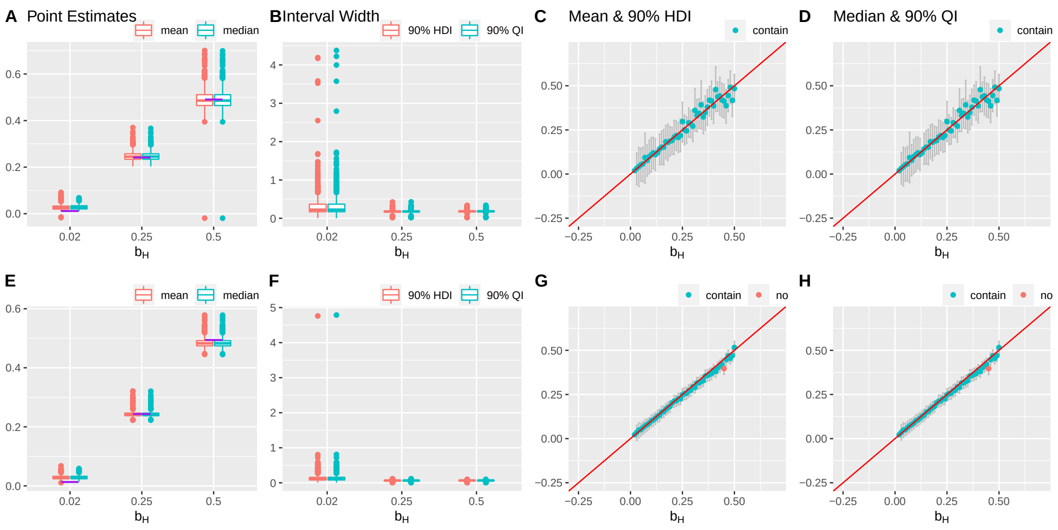

Figure S7: Results of a simulation setting: $n$ = 25,000, varying $b_H$, 1000 simulations. The first and second row presents the results for the 70 SNPs and 500 SNPs, respectively. (A, E) Box plots of the posterior median and the mean of $b_H$. The purple lines indicate the true values. (B, F) Box plots of the widths of 90% HDIs and QIs. (C, G) The posterior medians and the 90% HDIs of the 49 equally spaced values of $b_H$ between 0.02 and 0.5. (D, H) The posterior means and the 90% QIs of the 49 equally spaced values of $b_H$ between 0.02 and 0.5. Outliers are defined as the values more extreme than the third quartile + 1.5 * (the third quartile – the first quartile) or the first quartile – 1.5 * (the third quartile – the first quartile).

#### S3.4 Setting 4: different numbers of mediators

In Setting 4, we evaluate our method when there are ten or one known mediator(s) of the outcome trait. Based on the simulation results, the performance of our method does not change dramatically regardless of the number of known mediators, which shows the proposed method can deal with different numbers of known mediators in the model (Figures S8, S9, Table S1).

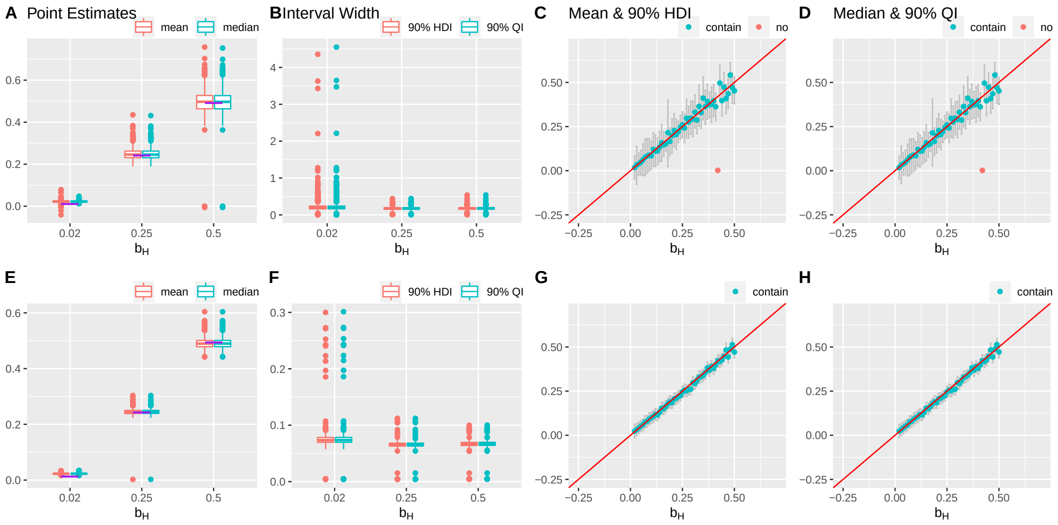

Figure S8: Results of a simulation setting: one known mediator, varying $b_H$, 1000 simulations. The first and second row presents the results for the 70 SNPs and 500 SNPs, respectively. (A, E) Box plots of the posterior median and the mean of $b_H$. The purple lines indicate the true values. (B, F) Box plots of the widths of 90% HDIs and QIs. (C, G) The posterior medians and the 90% HDIs of the 49 equally spaced values of $b_H$ between 0.02 and 0.5. (D, H) The posterior means and the 90% QIs of the 49 equally spaced values of $b_H$ between 0.02 and 0.5. Outliers are defined as the values more extreme than the third quartile + 1.5 * (the third quartile – the first quartile) or the first quartile – 1.5 * (the third quartile – the first quartile).

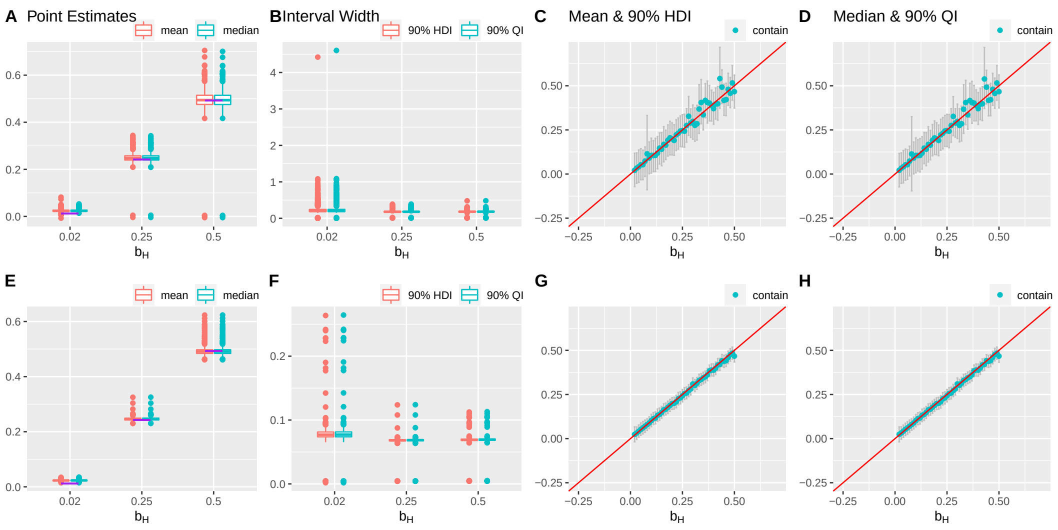

Figure S9: Results of a simulation setting: ten known mediators, varying $b_H$, 1000 simulations. The first and second row presents the results for the 70 SNPs and 500 SNPs, respectively. (A, E) Box plots of the posterior median and the mean of $b_H$. The purple lines indicate the true values. (B, F) Box plots of the widths of 90% HDIs and QIs. (C, G) The posterior medians and the 90% HDIs of the 49 equally spaced values of $b_H$ between 0.02 and 0.5. (D, H) The posterior means and the 90% QIs of the 49 equally spaced values of $b_H$ between 0.02 and 0.5. Outliers are defined as the values more extreme than the third quartile + 1.5 * (the third quartile – the first quartile) or the first quartile – 1.5 * (the third quartile – the first quartile).

#### S3.5 Setting 5: correlated mediators

In Setting 5, we allow causal relationships among the known mediators. We consider four cases. In the first case, there is a weak association between two of the known mediators ($M_1$ affects $M_2$ with an effect size of 0.3). In the second case, there is a relatively strong association between two of the known mediators ($M_1$ affects $M_2$ with an effect size of 0.9). In the third case, there are multiple weak associations among the known mediators ($M_1$ affects $M_2$ with an effect size of 0.3; $M_3$ affects $M_4$ with an effect size of 0.2). In the fourth case, there are multiple moderate associations among the known mediators ($M_1$ affects $M_2$ with an effect size of 0.5; $M_3$ affects $M_4$ with an effect size of 0.4). First, when the causal relationships are not adjusted in the mediation regressions during Step 1, only when there is a single weak association between two of the known mediators, are the results similar to the base case (Figure S10). In the other three cases, failure to adjust for the causal relations among the known mediators can lead to larger upward or downward biases in point estimates via either the median and the mean (Figures S11ACDEGH, S12ACDEGH, S13ACDEGH, Table S1). Also, both the HDI and the QI are less likely to capture the true value of $b_H$ under the 500 SNP case, and they are less likely to capture the true value of $b_H$ as the true value of $b_H$ increases (Figures S11BCDFGH, S12BCDFGH, S13BCDFGH, Table S1). Next, when the causal relationships among the known mediators are correctly adjusted for in the mediation regressions during Step 1, both the points estimators and the interval estimators perform similarly as the base case (Figures S10-13, Table S1). Thus, the results suggest that it is essential to adjust for the associations among the known mediators, which prior knowledge on the causal relationships among the known mediators may be required.

##### S3.5.1 Ignore the causal relationships

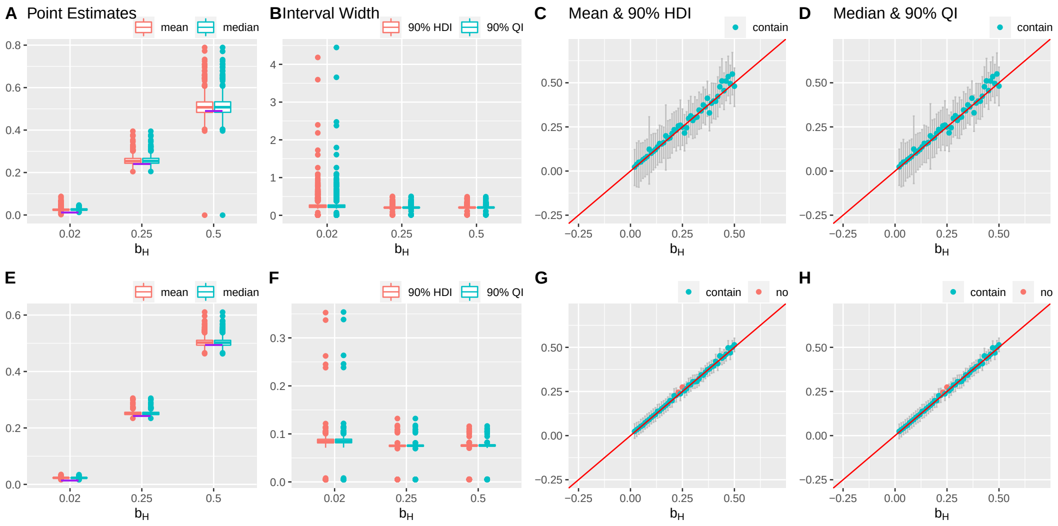

Figure S10: Results of a simulation setting: unadjusted, effect of $M_1$ on $M_2 = 0.3$, varying $b_H$, 1000 simulations. The first and second row presents the results for the 70 SNPs and 500 SNPs, respectively. (A, E) Box plots of the posterior median and the mean of $b_H$. The purple lines indicate the true values. (B, F) Box plots of the widths of 90% HDIs and QIs. (C, G) The posterior medians and the 90% HDIs of the 49 equally spaced values of $b_H$ between 0.02 and 0.5. (D, H) The posterior means and the 90% QIs of the 49 equally spaced values of $b_H$ between 0.02 and 0.5. Outliers are defined as the values more extreme than the third quartile + 1.5 * (the third quartile – the first quartile) or the first quartile – 1.5 * (the third quartile – the first quartile).

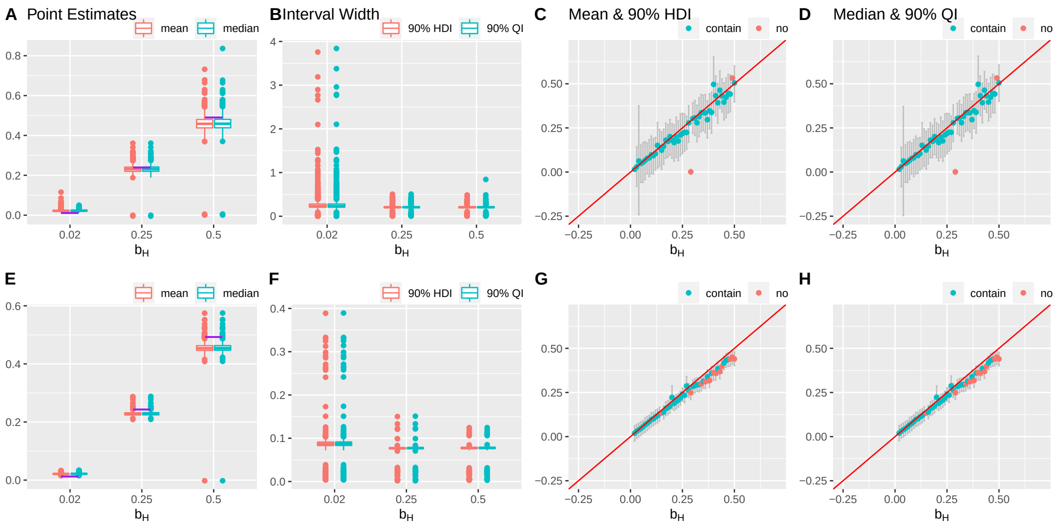

Figure S11: Results of a simulation setting: unadjusted, effect of $M_1$ on $M_2 = 0.9$, varying $b_H$, 1000 simulations. The first and second row presents the results for the 70 SNPs and 500 SNPs, respectively. (A, E) Box plots of the posterior median and the mean of $b_H$. The purple lines indicate the true values. (B, F) Box plots of the widths of 90% HDIs and QIs. (C, G) The posterior medians and the 90% HDIs of the 49 equally spaced values of $b_H$ between 0.02 and 0.5. (D, H) The posterior means and the 90% QIs of the 49 equally spaced values of $b_H$ between 0.02 and 0.5. Outliers are defined as the values more extreme than the third quartile + 1.5 * (the third quartile – the first quartile) or the first quartile – 1.5 * (the third quartile – the first quartile).

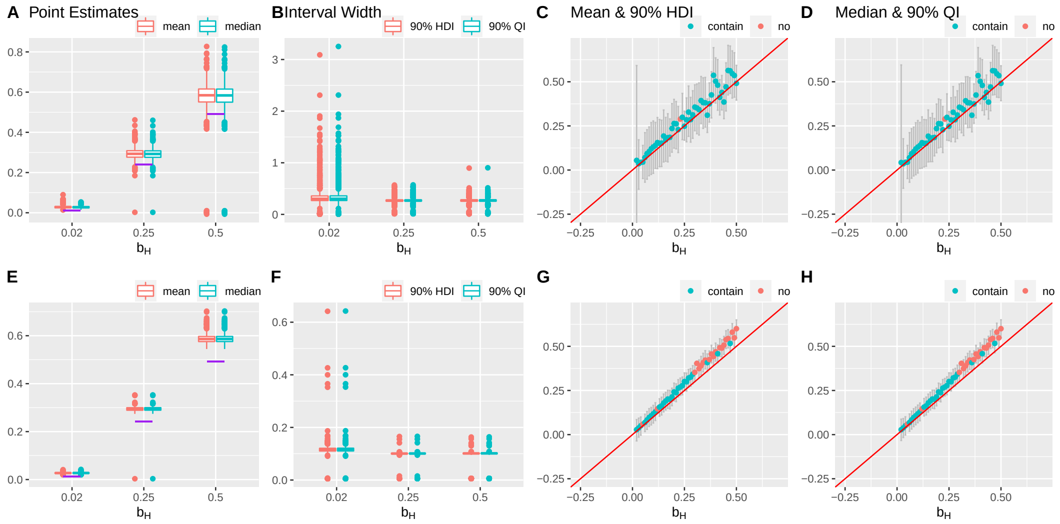

Figure S12: Results of a simulation setting: unadjusted, effect of $M_1$ on $M_2$ = 0.3, effect of $M_3$ on $M_4$ = 0.2, varying $b_H$, 1000 simulations. The first and second row presents the results for the 70 SNPs and 500 SNPs, respectively. (A, E) Box plots of the posterior median and the mean of $b_H$. The purple lines indicate the true values. (B, F) Box plots of the widths of 90% HDIs and QIs. (C, G) The posterior medians and the 90% HDIs of the 49 equally spaced values of $b_H$ between 0.02 and 0.5. (D, H) The posterior means and the 90% QIs of the 49 equally spaced values of $b_H$ between 0.02 and 0.5. Outliers are defined as the values more extreme than the third quartile + 1.5 * (the third quartile – the first quartile) or the first quartile – 1.5 * (the third quartile – the first quartile).

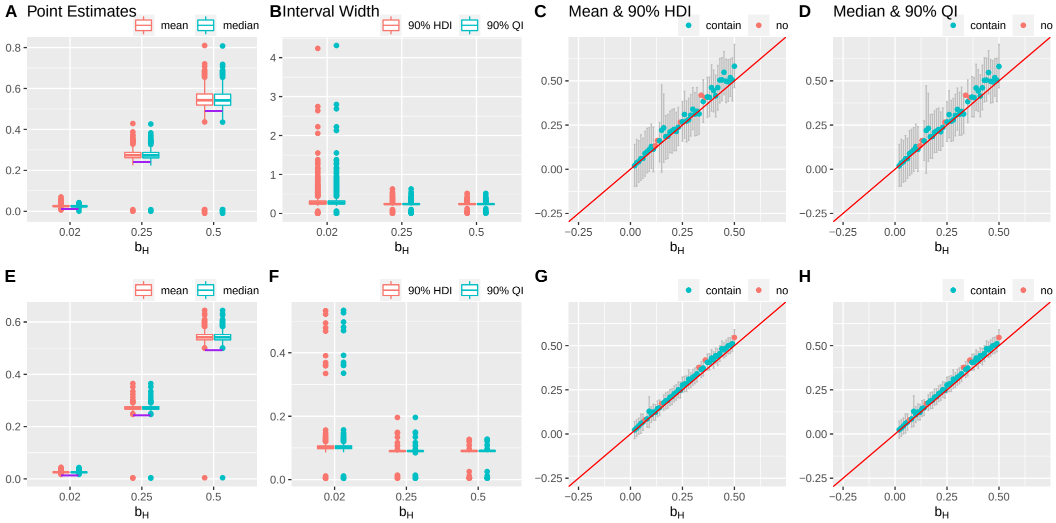

Figure S13: Results of a simulation setting: unadjusted, effect of $M_1$ on $M_2$ = 0.5, effect of $M_3$ on $M_4$ = 0.4, varying $b_H$, 1000 simulations. The first and second row presents the results for the 70 SNPs and 500 SNPs, respectively. (A, E) Box plots of the posterior median and the mean of $b_H$. The purple lines indicate the true values. (B, F) Box plots of the widths of 90% HDIs and QIs. (C, G) The posterior medians and the 90% HDIs of the 49 equally spaced values of $b_H$ between 0.02 and 0.5. (D, H) The posterior means and the 90% QIs of the 49 equally spaced values of $b_H$ between 0.02 and 0.5. Outliers are defined as the values more extreme than the third quartile + 1.5 * (the third quartile – the first quartile) or the first quartile – 1.5 * (the third quartile – the first quartile).

##### S3.5.2 Adjust for the causal relationships

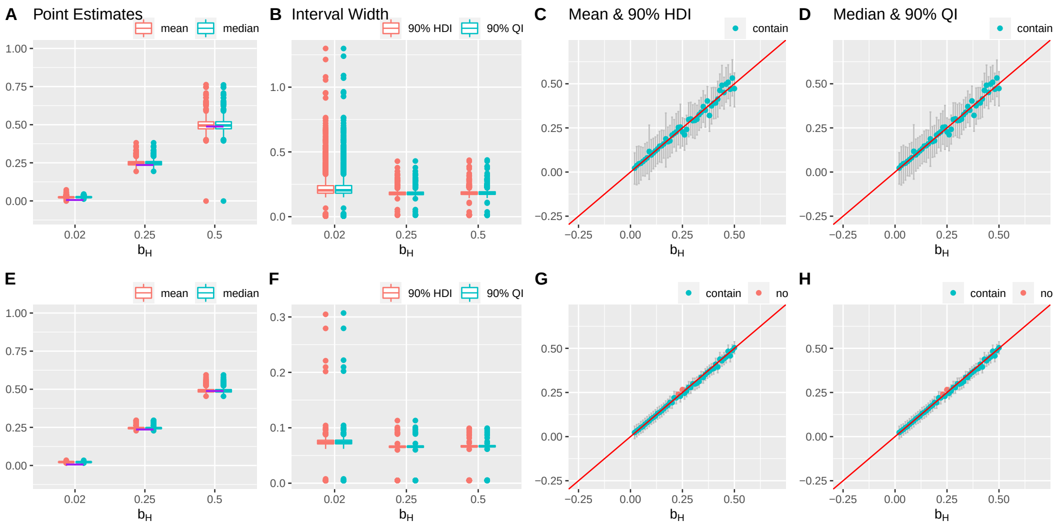

Figure S14: Results of a simulation setting: adjusted, effect of $M_1$ on $M_2 = 0.3$, varying $b_H$, 1000 simulations. The first and second row presents the results for the 70 SNPs and 500 SNPs, respectively. (A, E) Box plots of the posterior median and the mean of $b_H$. The purple lines indicate the true values. (B, F) Box plots of the widths of 90% HDIs and QIs. (C, G) The posterior medians and the 90% HDIs of the 49 equally spaced values of $b_H$ between 0.02 and 0.5. (D, H) The posterior means and the 90% QIs of the 49 equally spaced values of $b_H$ between 0.02 and 0.5. Outliers are defined as the values more extreme than the third quartile + 1.5 * (the third quartile – the first quartile) or the first quartile – 1.5 * (the third quartile – the first quartile).

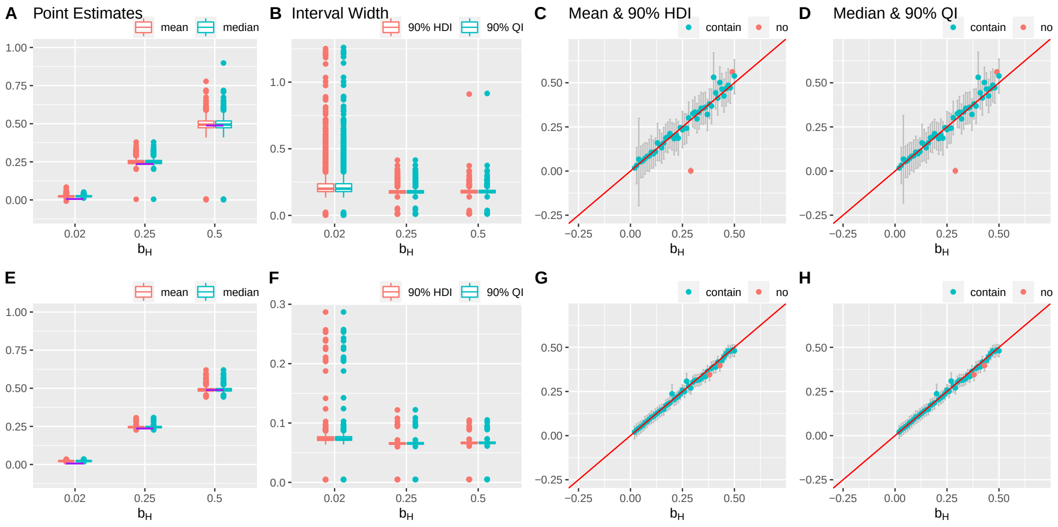

Figure S15: Results of a simulation setting: adjusted, effect of $M_1$ on $M_2 = 0.9$, varying $b_H$, 1000 simulations. The first and second row presents the results for the 70 SNPs and 500 SNPs, respectively. (A, E) Box plots of the posterior median and the mean of $b_H$. The purple lines indicate the true values. (B, F) Box plots of the widths of 90% HDIs and QIs. (C, G) The posterior medians and the 90% HDIs of the 49 equally spaced values of $b_H$ between 0.02 and 0.5. (D, H) The posterior means and the 90% QIs of the 49 equally spaced values of $b_H$ between 0.02 and 0.5. Outliers are defined as the values more extreme than the third quartile + 1.5 * (the third quartile – the first quartile) or the first quartile – 1.5 * (the third quartile – the first quartile).

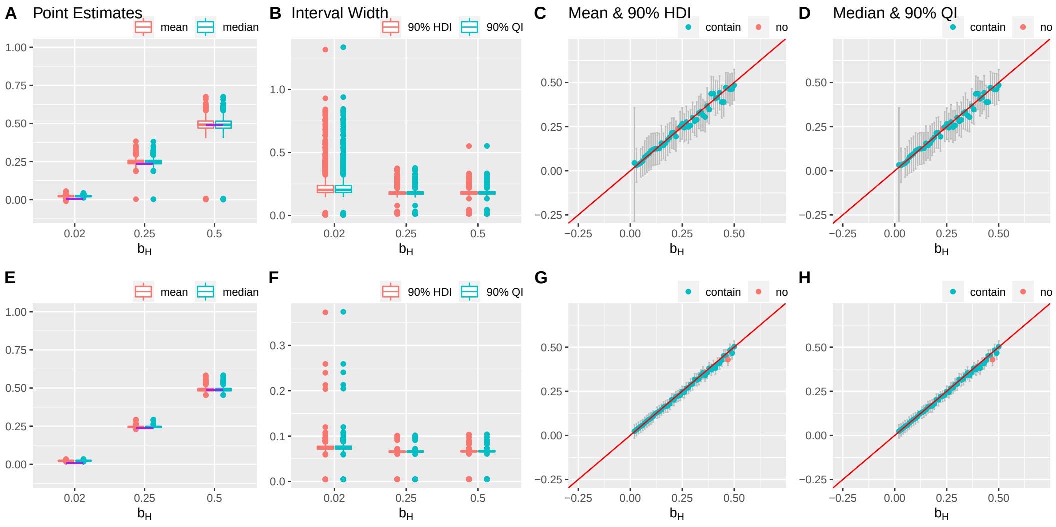

Figure S16: Results of a simulation setting: adjusted, effect of $M_1$ on $M_2 = 0.3$, effect of $M_3$ on $M_4 = 0.2$, varying $b_H$, 1000 simulations. The first and second row presents the results for the 70 SNPs and 500 SNPs, respectively. (A, E) Box plots of the posterior median and the mean of $b_H$. The purple lines indicate the true values. (B, F) Box plots of the widths of 90% HDIs and QIs. (C, G) The posterior medians and the 90% HDIs of the 49 equally spaced values of $b_H$ between 0.02 and 0.5. (D, H) The posterior means and the 90% QIs of the 49 equally spaced values of $b_H$ between 0.02 and 0.5. Outliers are defined as the values more extreme than the third quartile + 1.5 * (the third quartile – the first quartile) or the first quartile – 1.5 * (the third quartile – the first quartile).

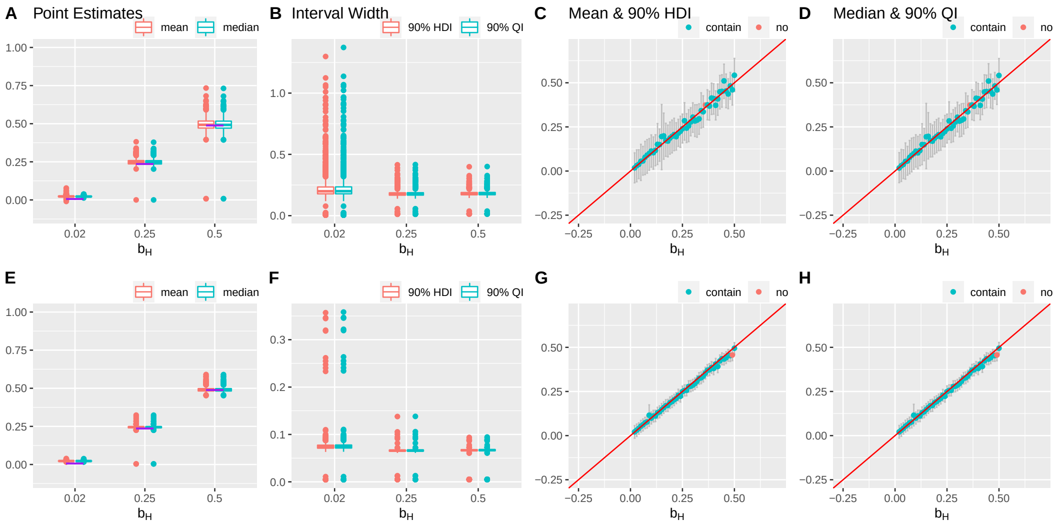

Figure S17: Results of a simulation setting: adjusted, effect of $M_1$ on $M_2 = 0.5$, effect of $M_3$ on $M_4 = 0.4$, varying $b_H$, 1000 simulations. The first and second row presents the results for the 70 SNPs and 500 SNPs, respectively. (A, E) Box plots of the posterior median and the mean of $b_H$. The purple lines indicate the true values. (B, F) Box plots of the widths of 90% HDIs and QIs. (C, G) The posterior medians and the 90% HDIs of the 49 equally spaced values of $b_H$ between 0.02 and 0.5. (D, H) The posterior means and the 90% QIs of the 49 equally spaced values of $b_H$ between 0.02 and 0.5. Outliers are defined as the values more extreme than the third quartile + 1.5 * (the third quartile – the first quartile) or the first quartile – 1.5 * (the third quartile – the first quartile).

#### S3.6 Setting 6: the three-level SNP effect structure

In this setting, the SNP effects on the mediators are generated from the three-level structure mentioned in Supplementary Material Section 1. The sum of the Level 1 and Level 2 standard deviation is fixed at 0.08. In the first case, the ratio between the Level 1 and Level 2 standard deviation is 1:3, which represents a scenario where the SNP effects on the hidden mediator are slightly different from the SNP effects on the known mediators. In the second case, the ratio between the Level 1 and Level 2 standard deviation is 1:1, which represents a scenario where the SNP effects on the hidden mediator are relatively different from the SNP effects on the known mediators. Although the median and the mean are almost unbiased, they are very imprecise with a large variation (Figure S18ACDEGH, S19ACDEGH). Increased variation is observed as the true value of $b_H$ increases, and the variation is bigger in the 1:1 case than in the 1:3 case. Also, the HDI and the QI are less likely to capture the true value of $b_H$ when the difference among the mediators increases (Figure S18BCDFGH, S19BCDFGH). Especially when there are 500 SNPs in the models, the interval estimators can lead to very wrong estimates when the true value of $b_H$ is large.

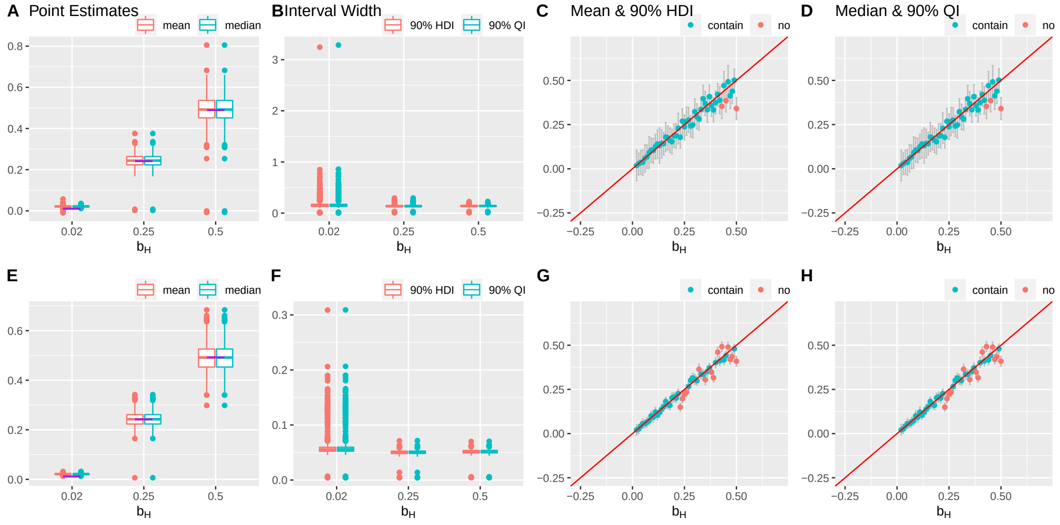

Figure S18: Results of a simulation setting: the first to second level standard deviation ratio = 1:3, varying $b_H$, 1000 simulations. The first and second row presents the results for the 70 SNPs and 500 SNPs, respectively. (A, E) Box plots of the posterior median and the mean of $b_H$. The purple lines indicate the true values. (B, F) Box plots of the widths of 90% HDIs and QIs. (C, G) The posterior medians and the 90% HDIs of the 49 equally spaced values of $b_H$ between 0.02 and 0.5. (D, H) The posterior means and the 90% QIs of the 49 equally spaced values of $b_H$ between 0.02 and 0.5. Outliers are defined as the values more extreme than the third quartile + 1.5 * (the third quartile – the first quartile) or the first quartile – 1.5 * (the third quartile – the first quartile).

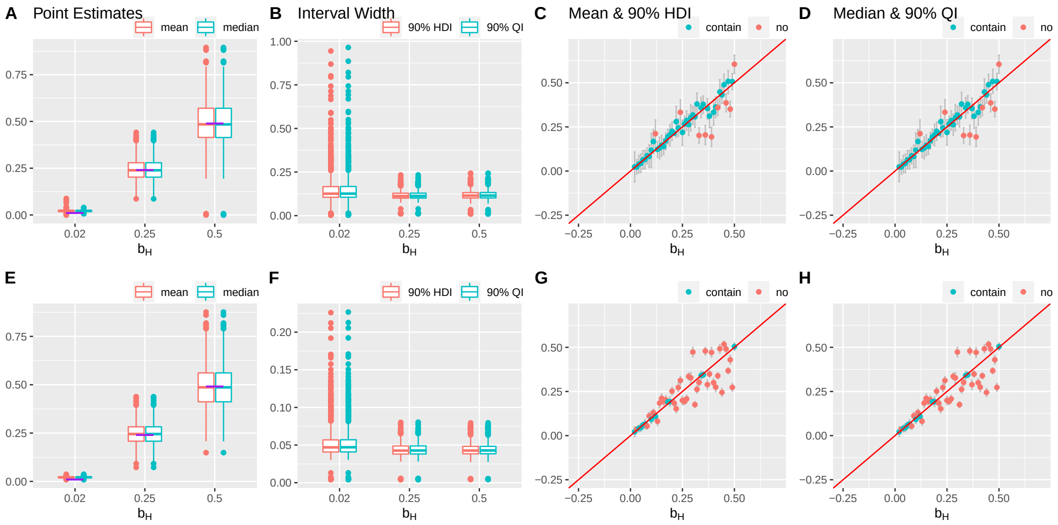

Figure S19: Results of a simulation setting: the first to second level standard deviation ratio = 1:1, varying $b_H$, 1000 simulations. The first and second row presents the results for the 70 SNPs and 500 SNPs, respectively. (A, E) Box plots of the posterior median and the mean of $b_H$. The purple lines indicate the true values. (B, F) Box plots of the widths of 90% HDIs and QIs. (C, G) The posterior medians and the 90% HDIs of the 49 equally spaced values of $b_H$ between 0.02 and 0.5. (D, H) The posterior means and the 90% QIs of the 49 equally spaced values of $b_H$ between 0.02 and 0.5. Outliers are defined as the values more extreme than the third quartile + 1.5 * (the third quartile – the first quartile) or the first quartile – 1.5 * (the third quartile – the first quartile).

#### S3.7 Setting 7: no hidden mediator

In Setting 7, $b_H$ is set to zero so there is no hidden mediator between the outcome trait the SNPs. Both the median and the mean are close to the true value of $b_H$, 0 (Figure S20ABEF). Despite some HDIs and QIs being relatively wide, the interval estimators capture the true value of $b_H$ well (Figure S20CDGH). The HDIs and QIs are wider relative to those in the other non-zero $b_H MJFZ$ settings. In addition, the capture rates of the HDI and the QI are close to 90%, and the type I error rates are close to 0.1, with the 500 SNP case's capture rate being closer to 90% and the type I error rate being closer to 0.1 (Table S2).

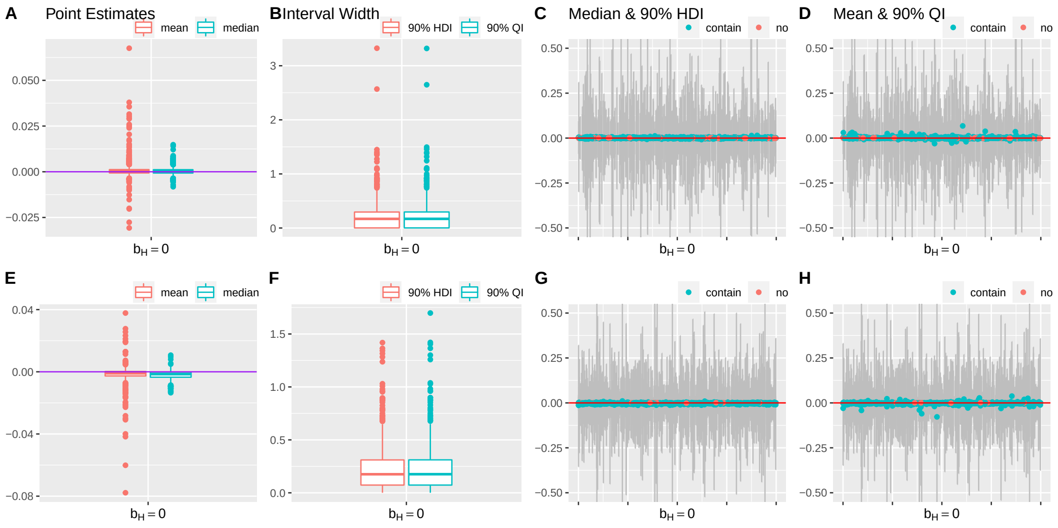

Figure S20: Results of a simulation setting: $b_H = 0$, 1000 simulations. The first and second row presents the results for the 70 SNPs and 500 SNPs, respectively. (A, E) Box plots of the posterior median and the mean of $b_H$. The purple lines indicate the true values. (B, F) Box plots of the widths of 90% HDIs and QIs. (C, G) The posterior medians and the 90% HDIs from the 1000 simulations. (D, H) The posterior means and the 90% QIs from the 1000 simulations. Outliers are defined as the values more extreme than the third quartile + 1.5 * (the third quartile – the first quartile) or the first quartile – 1.5 * (the third quartile – the first quartile).

#### S4 Proof: The distribution of $(a^*_{ij}|\Delta_{ij}=1)$ is $N(\mu_u,\ \sigma_u^2+\sigma_a^2+\sigma^2_{a^*_{ij}})$.

We sketch the proof to show that $\boldsymbol{a^*}$ and $\boldsymbol{c^*}$ follow a normal distribution. Therefore, fitting Gaussian mixture models (GMMs) on $\boldsymbol{a^*}$ and $\boldsymbol{c^*}$ is justified.

Based on Bayes rule, we have:

$$f(a^*_{ij},\ a_{ij},\ u_i|\Delta_{ij}=1) = f(a^*_{ij}|a_{ij},\ \Delta_{ij}=1)f(a_{ij}|u_i,\ \Delta_{ij}=1)f(u_i|\Delta_{ij}=1) \tag{S14}$$

Under the three-level SNP effect structure described in Supplementary Material Section 1, the second and third terms on the right-hand side of Equation S14 are both normal probability density functions. Because $\boldsymbol{a^*}$ are maximum likelihood estimates under linear regression models, the first term on the right-hand side of Equation S14 is also a normal probability density function.

Next, we integrate $u_i$ out of $f(a_{ij}|u_i,\ \Delta_{ij}=1)f(u_i|\Delta_{ij}=1)$ to obtain the distribution of $(a_{ij}|\Delta_{ij}=1)$. We start by writing out the normal probability density functions.

$$\begin{aligned} f(a_{ij}|u_i,\ \Delta_{ij}=1)f(u_i|\Delta_{ij}=1) &= \frac{1}{\sqrt{2\pi\sigma_a^2}}\exp\left(-\frac{(a_{ij}-u_i)^2}{2\sigma_a^2}\right)\frac{1}{\sqrt{2\pi\sigma_u^2}}\exp\left(-\frac{(u_i-\mu_u)^2}{2\sigma_u^2}\right) \\ &= \frac{1}{\sqrt{2\pi\sigma_a^2}}\exp\left(-\frac{a_{ij}^2-2a_{ij}u_i+u_i^2}{2\sigma_a^2}\right) \\ &\times \frac{1}{\sqrt{2\pi\sigma_u^2}}\exp\left(-\frac{u_i^2-2u_i\mu_u+\mu_u^2}{2\sigma_u^2}\right) \\ &= \frac{1}{\sqrt{2\pi\sigma_a^2}}\frac{1}{\sqrt{2\pi\sigma_u^2}}\exp\left(-\frac{a_{ij}^2}{2\sigma_a^2}-\frac{\mu_u^2}{2\sigma_u^2}\right) \\ &\times \exp\left\{-\frac{1}{2}\left[\left(\frac{1}{\sigma_a^2}+\frac{1}{\sigma_u^2}\right)u_i^2-2\left(\frac{a_{ij}}{\sigma_a^2}+\frac{\mu_u}{\sigma_u^2}\right)u_i\right]\right\} \end{aligned} \tag{S15}$$

Then we rewrite the exponential term in the last line of Equation S15 that involves $u_i$ as a normal probability density function.

$$\begin{aligned} &\exp\left\{-\frac{1}{2}\left[\left(\frac{1}{\sigma_a^2}+\frac{1}{\sigma_u^2}\right)u_i^2-2\left(\frac{a_{ij}}{\sigma_a^2}+\frac{\mu_u}{\sigma_u^2}\right)u_i\right]\right\} \\ &=\exp\left\{-\frac{1}{2}\left[\frac{u_i^2}{\frac{1}{\frac{1}{\sigma_a^2}+\frac{1}{\sigma_u^2}}}-\frac{2u_i\left(\frac{a_{ij}}{\sigma_a^2}+\frac{\mu}{\sigma_u^2}\right)\frac{1}{\frac{1}{\sigma_a^2}+\frac{1}{\sigma_u^2}}}{\frac{1}{\frac{1}{\sigma_a^2}+\frac{1}{\sigma_u^2}}}+\frac{\left(\frac{a_{ij}}{\sigma_a^2}+\frac{\mu}{\sigma_u^2}\right)^2\left(\frac{1}{\frac{1}{\sigma_a^2}+\frac{1}{\sigma_u^2}}\right)^2}{\frac{1}{\frac{1}{\sigma_a^2}+\frac{1}{\sigma_u^2}}}\right]\right\} \end{aligned} \tag{S16}$$

Note, Equation S16 is the kernel of the normal distribution $N\left[\left(\frac{a_{ij}}{\sigma_a^2}+\frac{\mu}{\sigma_u^2}\right)\left(\frac{1}{\frac{1}{\sigma_a^2}+\frac{1}{\sigma_u^2}}\right),\ \frac{1}{\frac{1}{\sigma_a^2}+\frac{1}{\sigma_u^2}}\right]$ with its probability density function being:

$$\frac{1}{\sqrt{2\pi\left(\frac{1}{\frac{1}{\sigma_a^2}+\frac{1}{\sigma_u^2}}\right)}}\exp\left\{-\frac{1}{2}\left[\frac{u_i^2}{\frac{1}{\frac{1}{\sigma_a^2}+\frac{1}{\sigma_u^2}}}-\frac{2u_i\left(\frac{a_{ij}}{\sigma_a^2}+\frac{\mu}{\sigma_u^2}\right)\frac{1}{\frac{1}{\sigma_a^2}+\frac{1}{\sigma_u^2}}}{\frac{1}{\frac{1}{\sigma_a^2}+\frac{1}{\sigma_u^2}}}+\frac{\left(\frac{a_{ij}}{\sigma_a^2}+\frac{\mu}{\sigma_u^2}\right)^2\left(\frac{1}{\frac{1}{\sigma_a^2}+\frac{1}{\sigma_u^2}}\right)^2}{\frac{1}{\frac{1}{\sigma_a^2}+\frac{1}{\sigma_u^2}}}\right]\right\}. \tag{S17}$$

Thus, integrating this normal probability density function with respect to $u_i$ gives 1. After integration, the remaining terms in Equation S14 does not involve $u_i$ and can be written as a normal probability density function as follow:

$$\begin{aligned}
&\frac{\sqrt{\frac{1}{\frac{1}{\sigma_a^2}+\frac{1}{\sigma_u^2}}}}{\sqrt{2\pi\sigma_a^2\sigma_u^2}} \exp\left(-\frac{a_{ij}^2}{2\sigma_a^2}-\frac{\mu_u^2}{2\sigma_u^2}\right) \exp\left\{-\frac{1}{2}\left[-\frac{\left(\frac{a_{ij}}{\sigma_a^2}+\frac{\mu}{\sigma_u^2}\right)^2\left(\frac{1}{\frac{1}{\sigma_a^2}+\frac{1}{\sigma_u^2}}\right)^2}{\frac{1}{\frac{1}{\sigma_a^2}+\frac{1}{\sigma_u^2}}}\right]\right\} \\
=&\frac{\sqrt{\frac{\sigma_a^2\sigma_u^2}{\sigma_a^2+\sigma_u^2}}}{\sqrt{2\pi\sigma_a^2\sigma_u^2}} \exp\left(-\frac{a_{ij}^2}{2\sigma_a^2}-\frac{\mu_u^2}{2\sigma_u^2}\right) \exp\left[\frac{1}{2}\left(\frac{a_{ij}}{\sigma_a^2}+\frac{\mu}{\sigma_u^2}\right)^2\left(\frac{\sigma_a^2\sigma_u^2}{\sigma_a^2+\sigma_u^2}\right)\right] \\
=&\frac{1}{\sqrt{2\pi\left(\sigma_a^2+\sigma_u^2\right)}} \exp\left(-\frac{a_{ij}^2}{2\sigma_a^2}-\frac{\mu_u^2}{2\sigma_u^2}\right) \exp\left[\frac{1}{2}\left(\frac{a_{ij}\sigma_u^2+\mu_u\sigma_a^2}{\sigma_a^2\sigma_u^2}\right)^2\left(\frac{\sigma_a^2\sigma_u^2}{\sigma_a^2+\sigma_u^2}\right)\right] \\
=&\frac{1}{\sqrt{2\pi\left(\sigma_a^2+\sigma_u^2\right)}} \exp\left[-\frac{\frac{a_{ij}^2}{\sigma_a^2}\left(\sigma_a^2+\sigma_u^2\right)+\frac{\mu_u^2}{2\sigma_u^2}\left(\sigma_a^2+\sigma_u^2\right)-\left(a_{ij}\sigma_u^2+\mu_u\sigma_a^2\right)^2\frac{1}{\sigma_a^2\sigma_u^2}}{2\left(\sigma_a^2+\sigma_u^2\right)}\right] \qquad \text{(S18)} \\
=&\frac{1}{\sqrt{2\pi\left(\sigma_a^2+\sigma_u^2\right)}} \exp\left[-\frac{a_{ij}^2+\frac{a_{ij}^2\sigma_u^2}{\sigma_a^2}+\frac{\mu_u^2\sigma_a^2}{\sigma_u^2}+\mu_u^2-\frac{a_{ij}^2\sigma_u^2}{\sigma_a^2}-2a_{ij}\mu_u-\frac{\mu_u^2\sigma_a^2}{\sigma_u^2}}{2\left(\sigma_a^2+\sigma_u^2\right)}\right] \\
=&\frac{1}{\sqrt{2\pi\left(\sigma_a^2+\sigma_u^2\right)}} \exp\left[-\frac{a_{ij}^2-2a_{ij}\mu_u+\mu_u^2}{2\left(\sigma_a^2+\sigma_u^2\right)}\right] \\
=&\frac{1}{\sqrt{2\pi\left(\sigma_a^2+\sigma_u^2\right)}} \exp\left[-\frac{\left(a_{ij}-\mu_u\right)^2}{2\left(\sigma_a^2+\sigma_u^2\right)}\right].
\end{aligned}$$

This is the probability density function of $N(\mu_u,\, \sigma_u^2+\sigma_a^2)$. Therefore, the distribution of $(a_{ij}|\Delta_{ij}=1)$ is $N(\mu_u,\, \sigma_u^2+\sigma_a^2)$. To summary, $u_i$ can be integrated out from Equation S14 to obtain

$$f(a_{ij}^*,\, a_{ij}|\Delta_{ij}=1)=f(a_{ij}^*|a_{ij},\, \Delta_{ij}=1)f(a_{ij}|\Delta_{ij}=1). \qquad \text{(S19)}$$

Next, similar steps can be applied to integrate $a_{ij}$ out of $f(a_{ij}^*,\, a_{ij}|\Delta_{ij}=1)$ to obtain $f(a_{ij}^*|\Delta_{ij}=1)$, and it can be found that $(a_{ij}^*|\Delta_{ij}=1)$ follows a normal distribution $N(\mu_u, \sigma_u^2+\sigma_a^2+\sigma_{a_{ij}^2})$. Thus, the regression estimated SNP effects on the mediators, $\boldsymbol{a^*}$, follow a normal distribution centered at the Level 1 mean $\mu_u$ under the three-level SNP effect structure. Following similar steps, it can be shown that the estimated direct effects between the outcome trait and its associated SNPs, $\boldsymbol{c^*}$, follow a normal distribution centered at $b_H\mu_u$. The detailed proof is not shown.

#### S5 Figure legends

1. Figure S1: The first two levels of the three-level SNP effect structure. A) The distribution of mediator-specific effect mean; B) The conditional distribution of SNP-specific effects.

2. Figure S2: Results of a simulation setting: $\pi_H = 0.3$, varying $b_H$, 1000 simulations. The first and second row presents the results for the 70 SNPs and 500 SNPs, respectively. (A, E) Box plots of the posterior median and the mean of $b_H$. The purple lines indicate the true values. (B, F) Box plots of the widths of 90% HDIs and QIs. (C, G) The posterior medians and the 90% HDIs of the 49 equally spaced values of $b_H$ between 0.02 and 0.5. (D, H) The posterior means and the 90% QIs of the 49 equally spaced values of $b_H$ between 0.02 and 0.5. Outliers are defined as the values more extreme than the third quartile + 1.5 * (the third quartile – the first quartile) or the first quartile – 1.5 * (the third quartile – the first quartile).

10. Figure S10: Results of a simulation setting: unadjusted, effect of $M_1$ on $M_2 = 0.3$, varying $b_H$, 1000 simulations. The first and second row presents the results for the 70 SNPs and 500 SNPs, respectively. (A, E) Box plots of the posterior median and the mean of $b_H$. The purple lines indicate the true values. (B, F) Box plots of the widths of 90% HDIs and QIs. (C, G) The

posterior medians and the 90% HDIs of the 49 equally spaced values of $b_H$ between 0.02 and 0.5. (D, H) The posterior means and the 90% QIs of the 49 equally spaced values of $b_H$ between 0.02 and 0.5. Outliers are defined as the values more extreme than the third quartile + 1.5 * (the third quartile – the first quartile) or the first quartile – 1.5 * (the third quartile – the first quartile).

14. Figure S14: Results of a simulation setting: adjusted, effect of $M_1$ on $M_2 = 0.3$, varying $b_H$, 1000 simulations. The first and second row presents the results for the 70 SNPs and 500 SNPs, respectively. (A, E) Box plots of the posterior median and the mean of $b_H$. The purple lines indicate the true values. (B, F) Box plots of the widths of 90% HDIs and QIs. (C, G) The

posterior medians and the 90% HDIs of the 49 equally spaced values of $b_H$ between 0.02 and 0.5. (D, H) The posterior means and the 90% QIs of the 49 equally spaced values of $b_H$ between 0.02 and 0.5. Outliers are defined as the values more extreme than the third quartile + 1.5 * (the third quartile – the first quartile) or the first quartile – 1.5 * (the third quartile – the first quartile).

18. Figure S18: Results of a simulation setting: the first to second level standard deviation ratio = 1:3, varying $b_H$, 1000 simulations. The first and second row presents the results for the 70 SNPs and 500 SNPs, respectively. (A, E) Box plots of the posterior median and the mean of $b_H$. The purple lines indicate the true values. (B, F) Box plots of the widths of 90% HDIs and QIs. (C,

G) The posterior medians and the 90% HDIs of the 49 equally spaced values of $b_H$ between 0.02 and 0.5. (D, H) The posterior means and the 90% QIs of the 49 equally spaced values of $b_H$ between 0.02 and 0.5. Outliers are defined as the values more extreme than the third quartile + 1.5 * (the third quartile – the first quartile) or the first quartile – 1.5 * (the third quartile – the first quartile).

19. Figure S19: Results of a simulation setting: the first to second level standard deviation ratio = 1:1, varying $b_H$, 1000 simulations. The first and second row presents the results for the 70 SNPs and 500 SNPs, respectively. (A, E) Box plots of the posterior median and the mean of $b_H$. The purple lines indicate the true values. (B, F) Box plots of the widths of 90% HDIs and QIs. (C, G) The posterior medians and the 90% HDIs of the 49 equally spaced values of $b_H$ between 0.02 and 0.5. (D, H) The posterior means and the 90% QIs of the 49 equally spaced values of $b_H$ between 0.02 and 0.5. Outliers are defined as the values more extreme than the third quartile + 1.5 * (the third quartile – the first quartile) or the first quartile – 1.5 * (the third quartile – the first quartile).

20. Figure S20: Results of a simulation setting: $b_H = 0$, 1000 simulations. The first and second row presents the results for the 70 SNPs and 500 SNPs, respectively. (A, E) Box plots of the posterior median and the mean of $b_H$. The purple lines indicate the true values. (B, F) Box plots of the widths of 90% HDIs and QIs. (C, G) The posterior medians and the 90% HDIs from the 1000 simulations. (D, H) The posterior means and the 90% QIs from the 1000 simulations. Outliers are defined as the values more extreme than the third quartile + 1.5 * (the third quartile – the first quartile) or the first quartile – 1.5 * (the third quartile – the first quartile).
